# AP-4 regulates neuronal lysosome composition, function, and transport via regulating export of critical lysosome receptor proteins at the trans-Golgi network

**DOI:** 10.1101/2021.06.25.449958

**Authors:** Piyali Majumder, Daisy Edmison, Catherine Rodger, Sruchi Patel, Evan Reid, Swetha Gowrishankar

**Author notes:** Correspondence to Swetha Gowrishankar.

## Abstract

The adaptor protein complex-4 or AP-4 is known to mediate autophagosome maturation through regulating sorting of transmembrane cargo such as ATG9A at the Golgi. There is a need to understand AP-4 function in neurons, as mutations in any of its four subunits cause a complex form of hereditary spastic paraplegia (HSP) with intellectual disability. While AP-4 has been implicated in regulating trafficking and distribution of cargo such as ATG9A and APP, little is known about its effect on neuronal lysosomal protein traffic, lysosome biogenesis and function. In this study, we demonstrate that in human iPSC-derived neurons AP-4 regulates lysosome composition, function and transport via regulating export of critical lysosomal receptors, including Sortilin 1, from the trans-Golgi network to endo-lysosomes. Additionally, loss of AP-4 causes endo-lysosomes to stall and build up in axonal swellings potentially through reduced recruitment of retrograde transport machinery to the organelle. These findings of axonal lysosome build-up are highly reminiscent of those observed in Alzheimer’s disease as well as in neurons modelling the most common form of HSP, caused by *spastin* mutations. Our findings implicate AP-4 as a critical regulator of neuronal lysosome biogenesis and altered lysosome function and axonal endo-lysosome transport as an underlying defect in AP-4 deficient HSP.

## Introduction

The adaptor protein complex-4 (AP-4) is a low abundance, ubiquitously expressed complex that belongs to the family of closely related hetero-tetrameric complexes (Dell’Angelica et al., 1999; Hirst et al., 1999) that are involved in sorting and trafficking of cargo in cells (Hirst et al., 2013). AP-4 complex consists of four subunits encoded by different genes: ε (*AP4E1*), μ (*AP4M1*), β (*AP4B1*) *and* σ (*AP4S1*). Mutations in any of these four genes result in a complex form of hereditary spastic paraplegia with intellectual disability, referred to as, AP-4 deficiency syndrome (Moreno-De-Luca et al., 2011; Verkerk et al., 2009; Abou Jamra et al., 2011; Tüysüz et al., 2014; Abdollahpour et al., 2015; Hardies et al., 2015; Ebrahimi-Fakhari et al., 2018).

Early *in vitro* and yeast-two hybrid studies that examined AP-4 interactions indicated that it binds to lysosomal proteins such as LAMP1 and LAMP2a as well as CD63 through interactions with YXXø motif (Bonifacino and Dell’Angelica, 1999). However, these interactions are weak, and loss of AP-4 function does not alter the distribution of these lysosomal proteins. Loss of AP-4 function does however strongly affect the intracellular traffic and distribution of the autophagy protein ATG9A (Mattera et al., 2017; De Pace et al., 2018; Davies et al., 2018). ATG9A is a multi-spanning transmembrane protein in the core autophagy machinery (Rubinsztein et al., 2012) that cycles between the trans-Golgi network (TGN) and peripheral organelles including endosomes and pre-autophagosomal structures, providing membrane for the growth of the latter (Reggiori et al., 2004; Young et al., 2006; Orsi et al., 2012; Imai et al., 2016). Loss of AP-4 in several cell types, including HeLa cells, MEFs and primary neurons results in strong retention of ATG9A in the TGN and reduced amounts of peripheral ATG9A leading to defects in autophagosomal maturation (De Pace et al., 2018; Davies et al., 2018; Mattera et al., 2017; Ivankovic et al., 2020).

While the involvement of AP-4 in regulating the autophagic pathway has received considerable attention, very little is known about how it affects lysosomal composition, trafficking and function. Studies have demonstrated that AP-4 ε KO mice exhibit axonal swellings in various regions of the brain and spinal cord (De Pace et al., 2018). Intriguingly, axonal swellings in the hippocampus and white matter tracts of the midbrain in AP-4 ε KO mice were enriched in the late endosomal and lysosomal protein, LAMP1 (Edmison et al., 2021; De Pace et al., 2018). This suggests that axonal endo-lysosome homeostasis is altered upon loss of AP-4.

Here, we examine the composition, distribution and function of lysosomes in AP-4 depleted human iPSC-derived neurons. We find that their neuronal lysosomes contain lower levels of certain lysosomal enzymes and exhibit compromised function. We show that loss of AP-4 dramatically alters distribution of Sortilin 1 (referred to as Sortilin here), a critical receptor involved in cargo transport between the TGN and endo-lysosomes (Nielsen et al., 2001). Lastly, we show that loss of AP-4 causes LAMP1-positive organelles as well as MAPK8IP3/JIP3, a neuronally enriched putative adaptor protein that regulates retrograde axonal transport, to build up in axonal swellings. Given AP-4 regulation of cargo sorting at the TGN, this could result from AP-4 loss altering TGN export of a regulator of axonal lysosome transport (and a potential interactor of MAPK8IP3), to the organelles. Thus, we demonstrate that the AP-4 complex is a key regulator of neuronal lysosome composition, function and transport. Our studies indicate that altered lysosome function and movement likely contribute to development of this form of complex HSP.

## Results

To determine the role of AP-4 complex in regulating neuronal lysosome function and traffic in human neurons, we made use of the CRISPRi - iPSC system (Wu et al., 2021; Tian et al., 2019; Fernandopulle et al., 2018; Wang et al., 2017). In iPSCs, the neurogenic transcription factor NGN2 is integrated under a doxycycline-responsive promoter at a safe harbor locus in the WTC11 iPSC line (Fernandopulle et al., 2018; Wang et al., 2017). This system allows simple and rapid generation of glutamatergic cortical neurons (i^3^Neurons) upon culture of the iPSCs in the presence of doxycycline for a brief period. These i^3^Neurons exhibit morphological and biochemical properties of neurons 14 days post-induction (Gowrishankar et al., 2021; Wang et al., 2017) and are electrically active after 21 days (Wang et al., 2017). We have employed a modified iPSC system, in which CRISPR-inhibition (CRISPRi) machinery is integrated into a safe harbor locus (Wu et al., 2021; Tian et al., 2019). In CRISPRi, an enzymatically dead Cas9 fused to a KRAB transcriptional repressor is targeted close to the transcriptional start site of the target gene by a single guide RNA (sgRNA), thereby inhibiting expression of the gene. This system has advantages over standard CRISPR-based knock-out systems, which include high specificity with low toxicity and strikingly few off-targets (Wu et al., 2021; Tian et al., 2019).

Depletion or loss of the ε subunit of AP-4 caused destabilization of the complex in cultured cells and in a mouse model (Ivankovic et al., 2020; Mattera et al., 2017) and should thus be effective in causing loss of AP-4 loss of function. We, therefore, transduced CRISPRi-iPSCs with either of two separate sgRNAs referred to as G2i or G4i (specific sequences of G2i and G4i in Supplemental Table S2), targeting the AP-4 ε transcriptional start site or a scrambled sequence (SC), then differentiated the resulting iPSC lines to neurons (*henceforth*: AP-4 and Control i^3^Neurons). After 21 days in culture, efficient depletion of AP-4 ε was observed in AP-4 CRISPRi i^3^Neurons relative to scrambled sgRNA control (Fig. 1 A, B).

**Figure 1.**
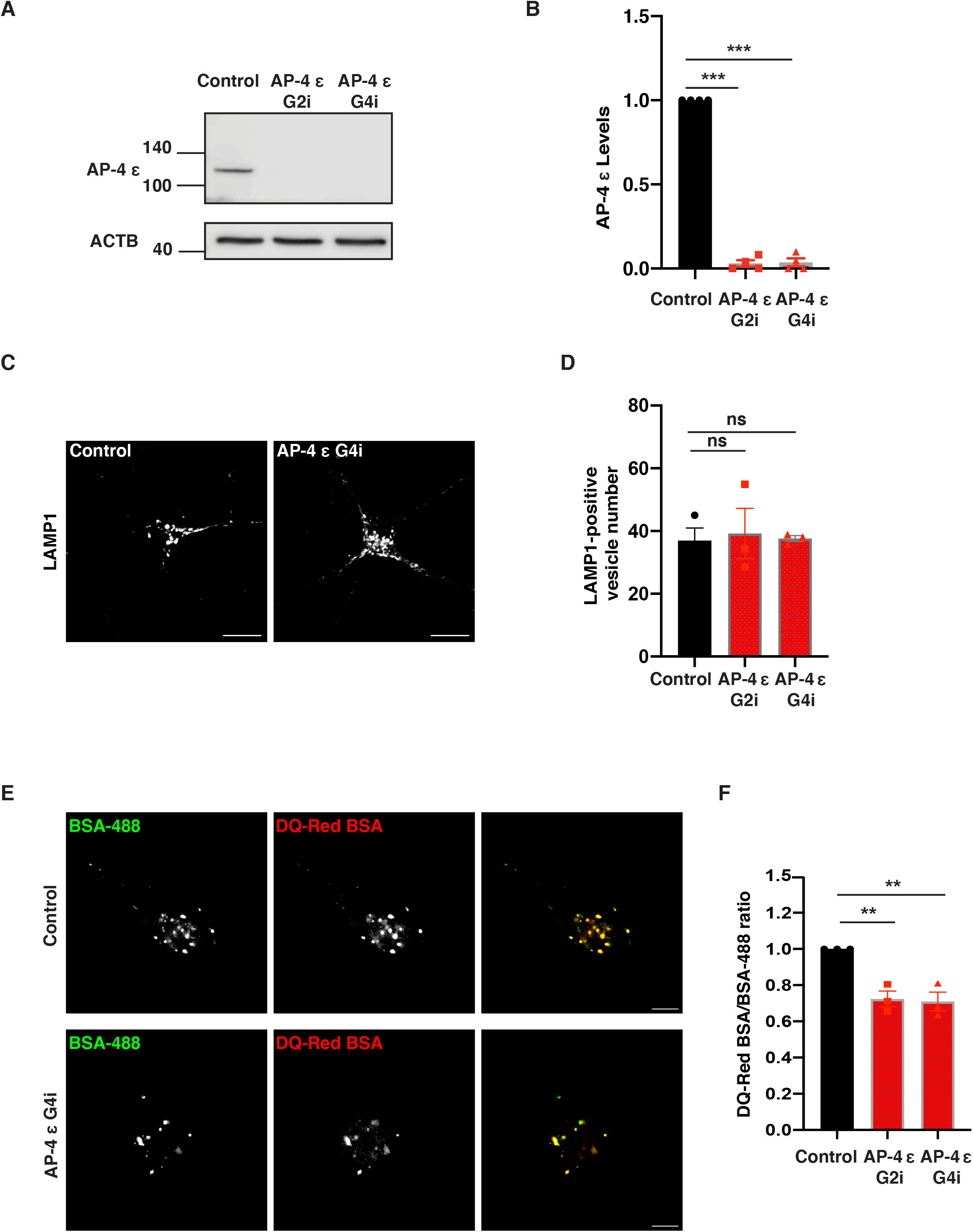
Loss of AP-4 affects lysosome function. (A and B) Immunoblotting reveals decreased levels of AP-4 protein in AP-4 i^3^Neurons compared to Control i^3^Neurons (21 DIV; ACTB used as loading Control; mean ± SEM from four independent experiments; ***, P < 0.001). (C) Control and AP-4 G4i i^3^Neurons (14 DIV) stained for LAMP1. (D) Quantification of number of LAMP1-positive vesicles in soma of AP-4 i^3^Neurons compared to Control i^3^Neurons (14 DIV; mean ± SEM from three independent experiments; >40 neurons per genotype; ns, not significant). (E) Images of DQ-Red BSA and A488 BSA fluorescence in Control and AP-4 G4i i^3^Neurons (14 DIV). Bar, 10 μm. (F) Quantification of lysosomal degradation (mean normalized DQ-Red BSA/BSA-488 ratio) in AP-4 i^3^Neurons compared to Control i^3^Neurons (14 DIV; mean ± SEM from three independent experiments; >40 neurons per genotype; **, P < 0.01).

We next examined the distribution of different lysosomal proteins in these i^3^Neurons at 14 days differentiation. Staining with LAMP1, a marker of late endosomes as well as degradative lysosomes (Lie et al., 2021; Gowrishankar et al., 2021; Cheng et al., 2018; Yap et al., 2018), revealed that there was not a significant difference in the number or distribution of LAMP1-positive vesicles in the cell bodies of AP-4 i^3^Neurons (Fig. 1 C, D; Fig. S1 A, B). AP-4 i^3^Neurons exhibited strong accumulation of ATG9A in the TGN (Fig. S2) when compared to Control i^3^Neurons, recapitulating the phenotype observed in other systems upon loss of AP-4 function (De Pace et al., 2018; Davies et al., 2018).

Since LAMP1 labels heterogenous organelles that include non-degradative vesicles as well as protease-rich degradative lysosomes (Cheng et al., 2018; Gowrishankar et al., 2015; Yap et al., 2018), we next examined lysosome function more directly in these i^3^Neurons using the DQ-Red BSA trafficking assay (Marwaha and Sharma, 2017) with some modifications for use in i^3^Neurons. This assay employs bovine serum albumin (BSA) that is heavily labeled with a BODIPY TR-X dye (DQ-Red BSA), that is self-quenched until the BSA is degraded in lysosomes to smaller protein fragments resulting is de-quenching of the isolated dye molecules and increased fluorescence, pulsed along with A488 BSA whose fluorescence serves as a control for endocytic uptake. Lysosomal degradative efficiency, as read out by fluorescence of DQ-Red BSA normalized to fluorescence of A488 BSA for each cell was found to decrease by 25-30% in AP-4 i^3^Neurons compared to the Control i^3^Neurons (Fig. 1 E, F; Fig. S1 C, D).

In view of the reduced efficiency of lysosome proteolytic function in AP-4 i^3^Neurons, we next examined the maturation and distribution of lysosomal proteases in these cells. We found that the maturation of the cysteine protease, Cathepsin L was severely affected in AP-4 i^3^Neurons, with the amount of mature processed Cathepsin L in AP-4 i^3^Neurons about 50% of that in Control i^3^Neurons (Fig. 2 A, B). We also found that the number of Cathepsin L-positive organelles in the soma of AP-4 i^3^Neurons was about 50% of that in Control i^3^Neurons (Fig. 2 C, D; Fig. S3 A, B). Likewise, there was a strong reduction in the number of PPT-1 enzyme-containing lysosomes (Fig. 2 E, F; Fig. S3 C, D), as well as some reduction in total cellular PPT-1 enzyme levels (Fig. S3 E, F). While the number of PPT-1 and Cathepsin L-positive vesicles are reduced, they each colocalize extensively with LAMP1 indicating they are lysosomal (Fig. S4; Fig. S5). Given AP-4’s previously established role in regulating cargo sorting and export out the Golgi, we hypothesized that the reduction in levels of proteases at lysosomes likely resulted from perturbed Golgi to endo-lysosome transport of these cargo. While Cathepsin L and PPT-1 were affected, another protease, Cathepsin B, was not affected (Fig S6). AP-4 i^3^Neurons have similar number of Cathepsin B positive lysosomes when compared with Control i^3^Neurons (Fig. S6 A-C), suggesting that AP-4 specifically controls trafficking of a subset of lysosomal proteases.

**Figure 2.**
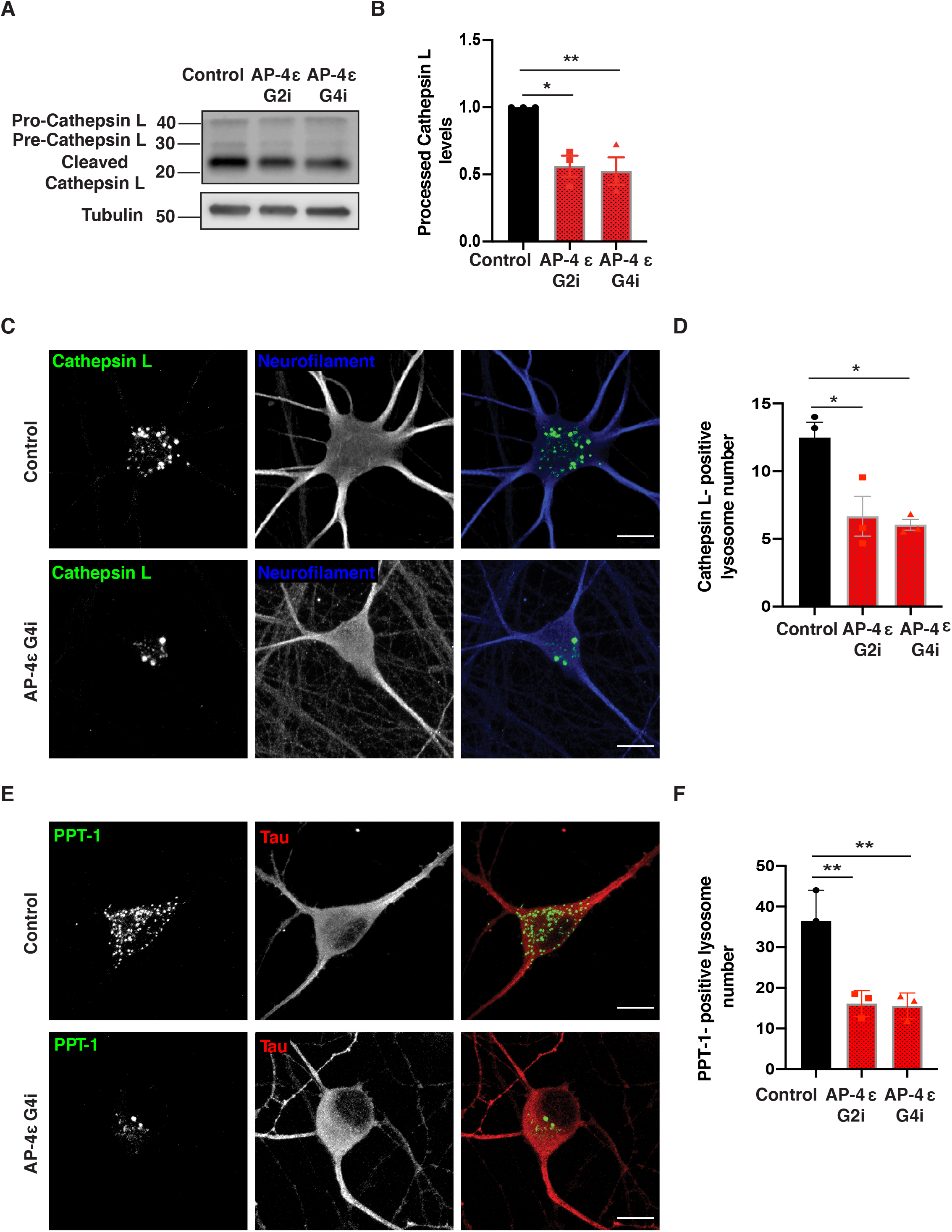
Loss of AP-4 affects neuronal lysosomal composition. (A and B) Immunoblotting reveals decreased levels of Cleaved/Pro-Cathepsin L protein in AP-4 i^3^Neurons compared to Control i^3^Neurons (21 DIV; α-Tubulin used as loading Control; mean ± SEM from three independent experiments; *, P < 0.05; **, P < 0.01). (C) Control and AP-4 G4i i^3^Neurons (14 DIV) stained for Cathepsin L (green) and neurofilament (blue), showing a reduced number of Cathepsin L-enriched lysosomes in cell bodies of AP-4 i^3^Neurons. Bar, 10 μm. (D) Quantification of Cathepsin L-positive vesicles in AP-4 i^3^Neurons compared to Control i^3^Neurons (14 DIV; mean ± SEM from three independent experiments; 25-30 neurons per genotype; *, P < 0.05). (E) Control and AP-4 G4i i^3^Neurons (14 DIV) stained for PPT-1 (green) and Tau (red). Bar, 10 μm. (F) Quantification of PPT-1 positive vesicles in AP-4 i^3^Neurons compared to Control i^3^Neurons (14 DIV; mean ± SEM from three independent experiments; >30 neurons per genotype; **, P < 0.01).

The trafficking of lysosomal proteases from the Golgi to endo-lysosomes is dependent on their interaction with receptors such as Mannose-6-phosphate receptor (M6PR) and Sortilin (Koster and Yoshii, 2019; Qian et al., 2008; Braulke and Bonifacino, 2009). Examination of Sortilin revealed a dramatic change in its intracellular distribution. While there was a strong build-up of Sortilin at the TGN area in AP-4 i^3^Neurons (Fig. 3 A, B; Fig. S7 A, B; Fig. S8), Sortilin signal in Control i^3^Neurons appears much dimmer with relatively few, faint puncta visible (Fig. S7 B). We confirmed that the Sortilin accumulation in the soma of AP-4 i^3^Neurons is in the TGN as it colocalized with TGN46 (Fig. S8). This retention of Sortilin at TGN in AP-4 i^3^Neurons suggests that Sortilin export from TGN is AP-4 dependent. Reduction in PPT-1 and Cathepsin L-containing lysosomes would also suggest that the delivery of these lysosomal enzymes may be Sortilin-dependent in neurons.

**Figure 3.**
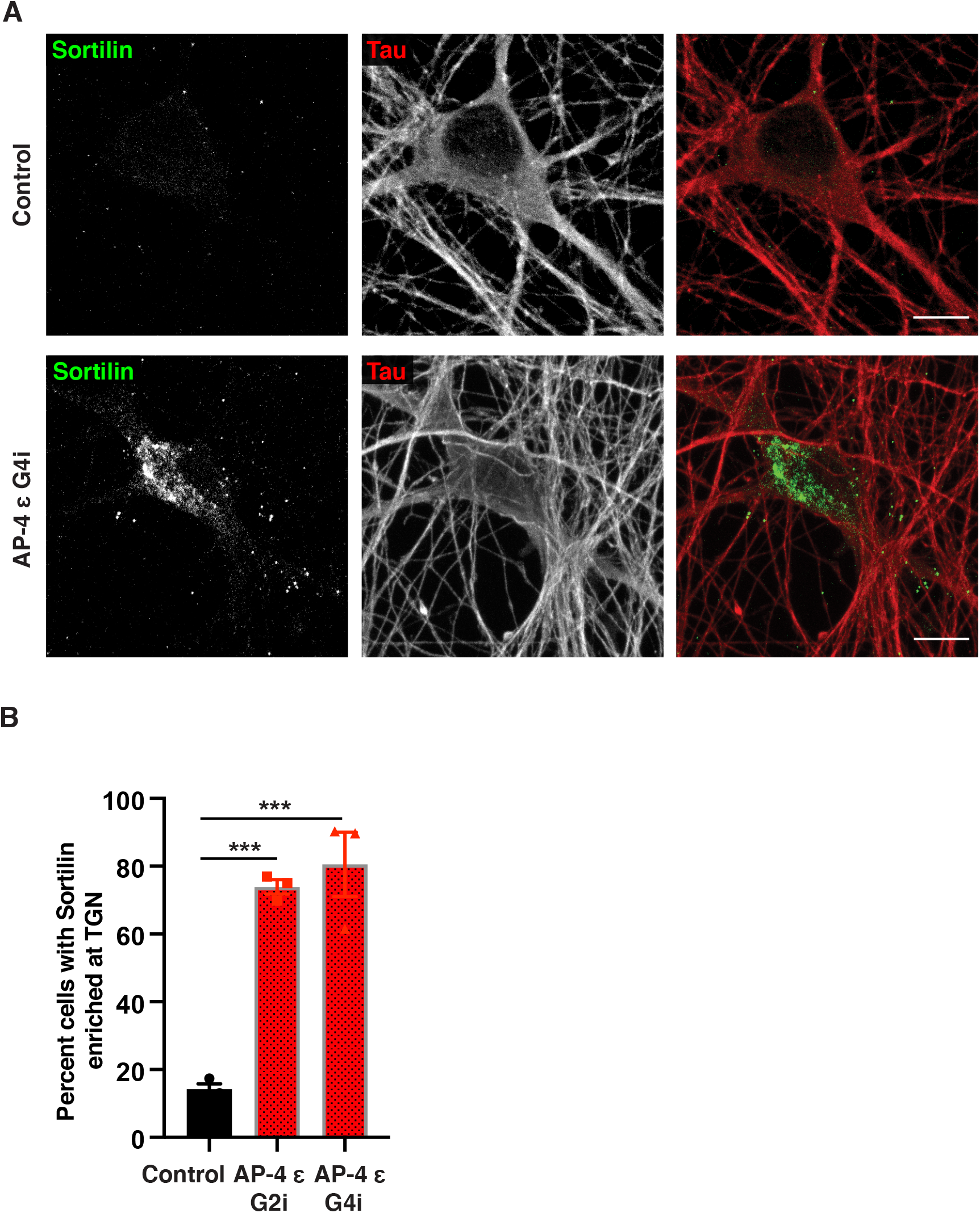
Sortilin distribution is dramatically altered in AP-4 i^3^Neurons. (A) Representative images of Control and AP-4 G4i i^3^Neurons (42 DIV) stained for Sortilin (green) and Tau (red), showing Sortilin is strongly accumulated in the TGN area in soma of AP-4 i^3^Neurons. Bar, 10 μm. (B) Quantification of the percentage of AP-4 i^3^Neurons with Sortilin accumulation in the TGN area compared to Control i^3^Neurons (42 DIV; mean ± SEM from three independent experiments; >100 neurons per genotype; ***, P < 0.001).

Having observed changes in lysosome function and composition in the soma, we next focused on LAMP1 distribution in axons. In Control i^3^Neurons, consistent with previous reports (Gowrishankar et al., 2021), most LAMP1 vesicles are largely concentrated in the soma with relatively fewer vesicles observed in neurites (Fig. 4 A, B). In contrast, we found that AP-4 i^3^Neurons robustly developed Tau-positive axonal dystrophies with age (at DIV 42) and these axonal swellings were filled with LAMP1-positive vesicles (Fig. 4 A, B; Fig. S9 A, D; white arrows). We confirmed that these LAMP1 accumulations were axonal as they were negative for the dendritic marker, MAP2B (Fig. 4 C; Fig. S9 B). Airyscan imaging of such axonal swellings revealed the presence of numerous ring-like and punctate LAMP1-positive vesicles in them (Video 1). Since axonal LAMP1-vesicles can be heterogenous, composed of both biosynthetic LAMP1-carriers as well as maturing endo-lysosomes (Gowrishankar et al., 2021; Lie et al., 2021), we used LysoTracker staining to determine if the vesicles that built up were acidic. Indeed, large, LysoTracker-positive (acidic) vesicles moved retrogradely towards or were stalled in axonal swellings in AP-4 i^3^Neurons (Video 2; Fig. S9 C). Since LAMP1-positive acidic vesicles exhibit a predominantly retrograde movement in axons (Gowrishankar et al., 2021; Lie et al., 2021), we examined the distribution and levels of JIP3/MAPK8IP3, a neuronally enriched scaffolding protein that potentially links axonal endo-lysosomes and motors and regulates their retrograde axonal transport (Gowrishankar et al., 2021; Rafiq et al., 2022). Interestingly, JIP3 protein level is significantly increased in AP-4 i^3^Neurons compared to Control i^3^Neurons (Fig. 5 A, B). When we examined distribution of endogenous JIP3 in the i^3^Neurons, we found that the AP-4 i^3^Neurons exhibited multiple JIP3-positive axonal swellings, while there was almost no JIP3 build up in Control i^3^Neurons (Fig. 5 C, D; Fig. S10 A, B). Thus, loss of AP-4 in neurons causes LAMP1-positive vesicles as well JIP3/MAPK8IP3 to build up in axonal dystrophies.

**Figure 4.**
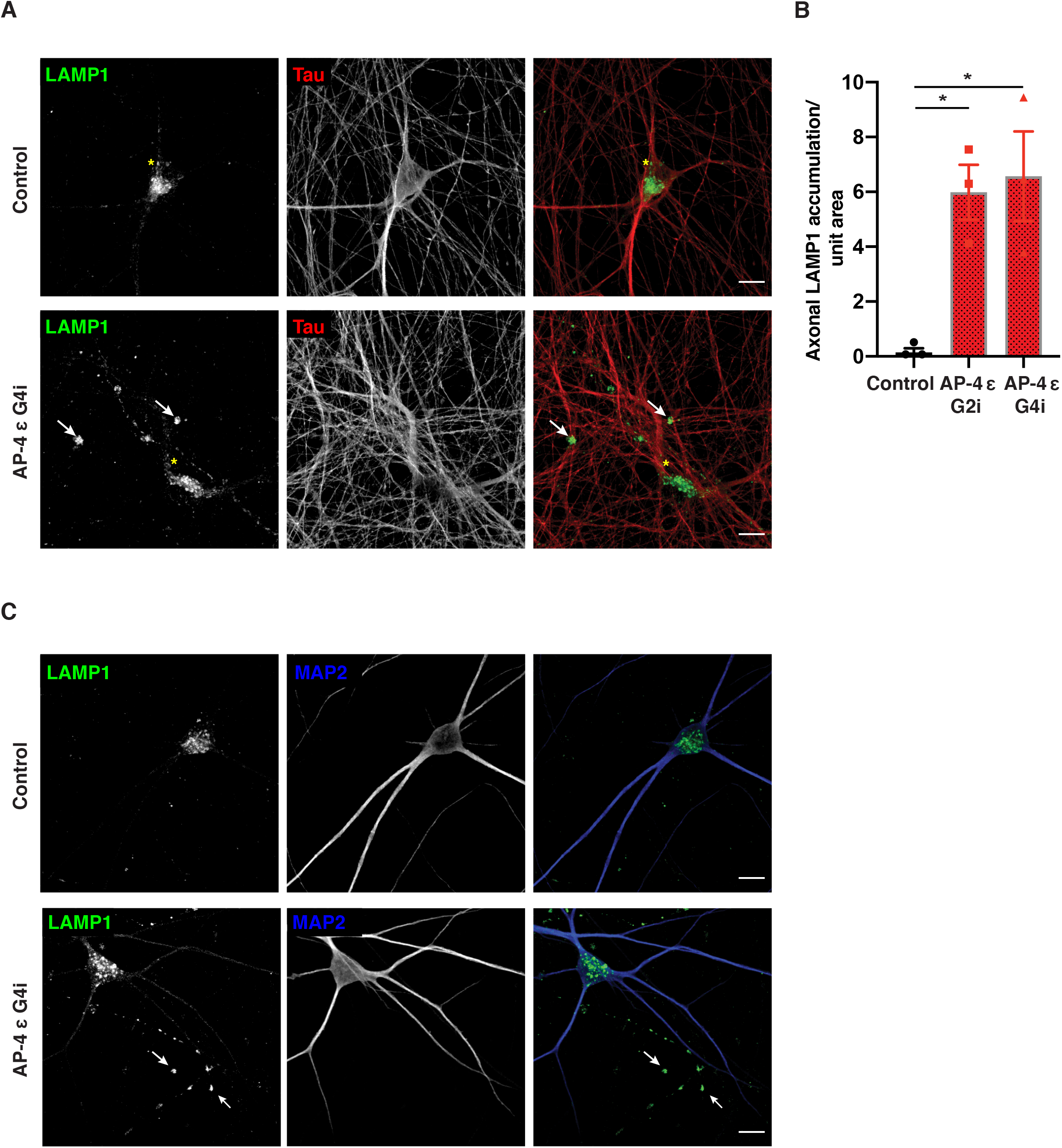
Loss of AP-4 in i^3^Neurons results in axonal lysosome build-up. (A) Control and AP-4 G4i i^3^Neurons (42 DIV) stained for LAMP1 (green) and Tau (red). White arrows highlight LAMP1 accumulation in the neurites of AP-4 i^3^Neurons. Yellow asterisks indicate the LAMP1-positive vesicles in the cell body. Bar, 20 μm. (B) Quantification of axonal LAMP-positive swellings per unit area in AP-4 i^3^Neurons and Control i^3^Neurons (42 DIV; mean ± SEM from three independent experiments; > 20 areas imaged at random per genotype; *, P < 0.05). (C) Representative images of Control and AP-4 G4i i^3^Neurons (42 DIV) stained for LAMP1 (green) and MAP2 (blue). Arrowheads highlight LAMP1 accumulation in the MAP2-negative neurites (axons) of AP-4 i^3^Neurons. Bar, 20 μm.

**Figure 5.**
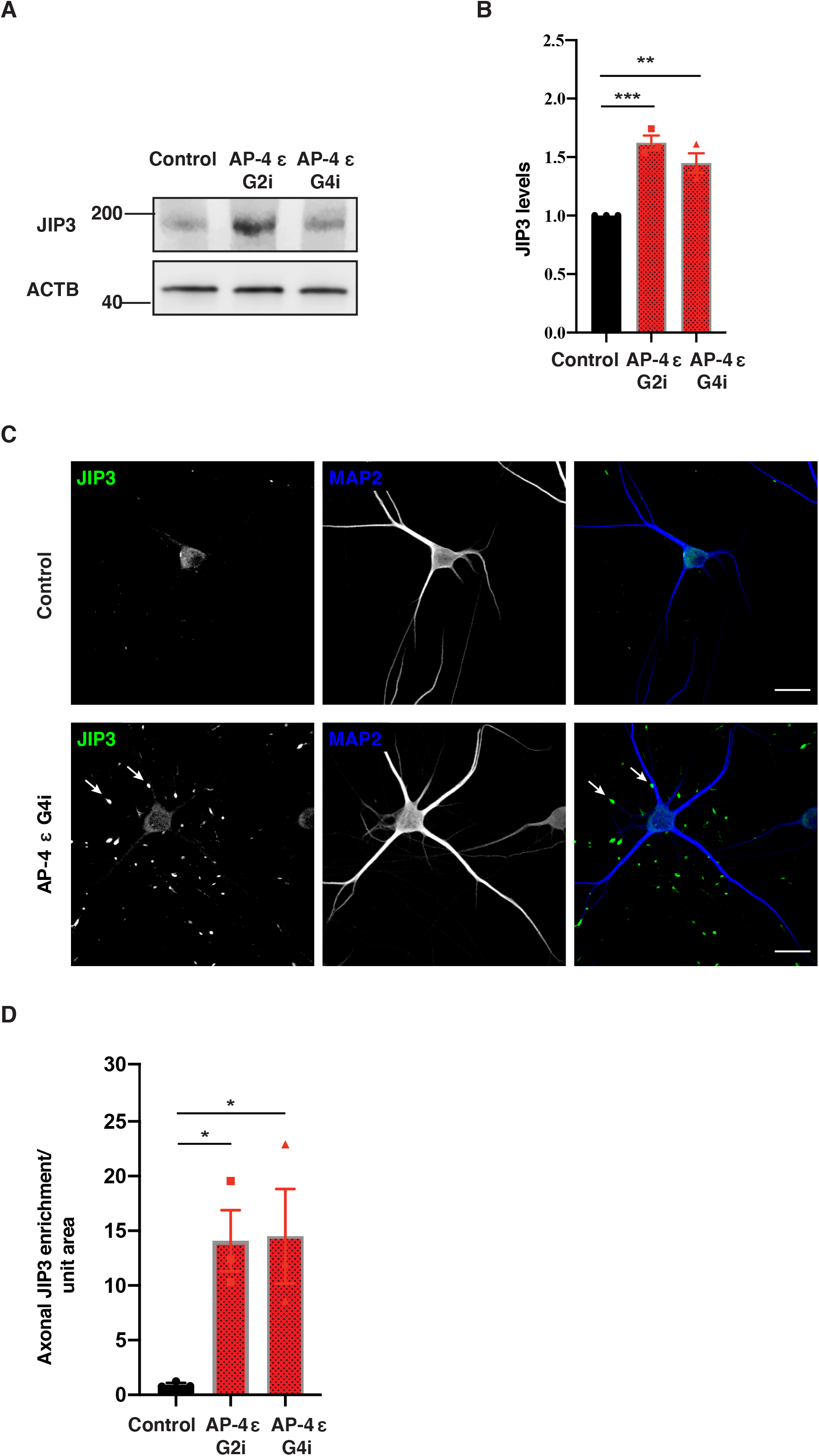
Loss of AP-4 in i^3^Neurons results in accumulation of JIP3 in axonal swellings. (A and B) Immunoblotting reveals increased levels of JIP3 in AP-4 i^3^Neurons compared to Control i^3^Neurons (21 DIV; ACTB used as loading Control; mean ± SEM from three independent experiments; **, P < 0.01; ***, P < 0.001). (C) Representative images of Control and AP-4 G4i i^3^Neurons (28 DIV) stained for JIP3 (green) and MAP2 (blue). Arrowheads highlight JIP3 accumulation in the axons of AP-4 G4i i^3^Neurons. Bar, 20 μm. (D) Quantification of axonal JIP3-positive swellings per unit area in AP-4 i^3^Neurons compared to Control i^3^Neurons (28 DIV; mean ± SEM from three independent experiments; > 20 areas chosen at random per genotype; *, P < 0.05).

## Discussion

Loss of AP-4 function leads to “AP-4 deficiency syndrome” which is also a form of complex HSP with intellectual disability. While AP-4 is ubiquitously expressed and its loss affects different systems, the nervous system is particularly adversely impacted, consistent with the idea that neurons exhibit greater vulnerability to impaired protein trafficking and lysosome dysfunction. A key aspect to understanding disease pathology arising from loss of function of this coat protein complex is to identify the different cargoes whose transport it regulates in neurons. While studies have demonstrated that AP-4 regulates traffic of proteins linked to the autophagy pathway, relatively little is known about how it may affect the closely related lysosomal pathway, a central player in maintaining protein and organelle homeostasis in neurons. In this study, we present evidence that neuronal lysosome composition and function as well as axonal movement and distribution of these organelles are affected by loss of AP-4.

### AP-4 CRISPRi i^3^Neurons as a model for examining pathology

The AP-4 depleted i^3^Neurons based on CRISPRi technology (Tian et al., 2019) are a good model system for studying alterations in cargo traffic in human neurons upon AP-4 loss. Apart from a robust depletion of the AP-4 ε subunit, we demonstrate that these i^3^Neurons phenocopy the strong TGN-retention of ATG9A as well as increased abundance of ATG9A (Fig. S2) observed in other neuronal and non-neuronal model systems (De Pace et al., 2018; Davies et al., 2018). We have also identified changes in the neuron-enriched protein JIP3/MAPK8IP3, alterations to which have been linked to a neurodevelopmental disorder (Platzer et al., 2019; Iwasawa et al., 2019). This reiterates the importance of exploring cellular changes in response to loss of AP-4 function in a neuronal model.

### AP-4 and neuronal lysosome composition and function

Studies in non-neuronal cells indicate that AP-4 can interact with sorting motifs in tails of receptor proteins such as M6PR, Furin and LDL receptor (Dennes et al., 2002; Nilsson et al., 2008; Jacobsen et al., 2001; Braulke and Bonifacino, 2009; Nielsen et al., 2001). Here, we have demonstrated that loss of AP-4 causes reduced abundance of certain lysosomal enzymes in LAMP1-positive organelles. Considering the role of AP-4 in sorting and export of other cargo from the TGN, this probably results from reduced biosynthetic delivery of these enzymes from the TGN to endo-lysosomes. Since many lysosomal enzymes are dependent on M6PR and/or Sortilin to get sorted at the TGN and trafficked to endo-lysosomes (Saftig and Klumperman, 2009; Braulke and Bonifacino, 2009), we examined the distribution of these receptors in AP-4 i^3^Neurons. We found that Sortilin is strongly retained in the TGN in AP-4 i^3^Neurons (Fig. 6; magnified portion of schematic of soma in AP-4 i^3^Neurons), in contrast to the more vesicular distribution in Control i^3^Neurons.

**Figure 6.**
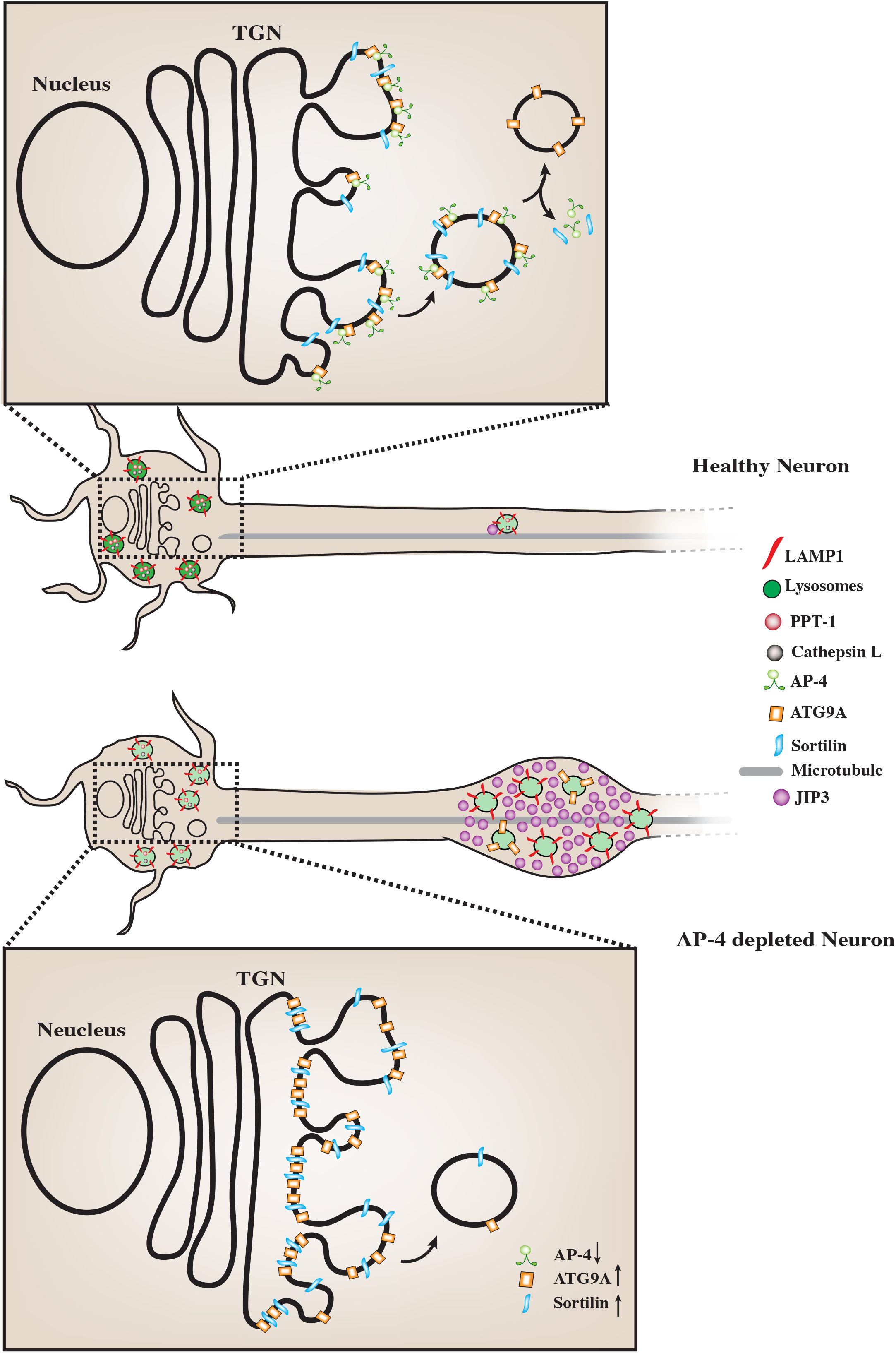
AP-4 regulates neuronal lysosome composition, function and transport. Schematic representation of the effect of AP-4 loss on neuronal lysosome trafficking and function. In AP-4 i^3^Neurons there are lysosomes in the soma that contain reduced levels of certain lysosomal enzymes, such as PPT-1 and Cathepsin L. Magnified insets of the TGN area in the soma of the neurons show molecular detail of cargo potentially regulated by AP-4 action. Upon loss of AP-4 complex function, concentration and transport of Sortilin out of the TGN is affected, when compared to healthy Control i^3^Neurons. Thus, strong Sortilin accumulation in the TGN area is observed in the AP-4 i^3^Neurons. ATG9A, previously well-established cargo whose sorting out of the TGN is dependent on AP-4 complex function, shows a similar strong accumulation in TGN area, validating these CRISPRi-based AP-4 i^3^Neurons as a model for studying neuronal membrane trafficking events regulated by AP-4 complex. Also, unlike healthy Control i^3^Neurons, where few LAMP1-positive vesicles are observed in axons, AP-4 loss causes axonal lysosomes (LAMP1-positive vesicles) to build up in dystrophies/swellings. Additionally, there is increased accumulation of JIP3/MAPK8IP3, a regulator of retrograde axonal lysosome transport, in these swellings. We propose that the accumulation of cytosolic JIP3 in the axonal swellings in AP-4 i^3^Neurons could also be due to absence/reduced levels of these interacting partners on lysosomes, whose sorting from TGN to the organelle is AP-4 dependent, as with ATG9A and potentially Sortilin.

### AP-4 function and links to APP processing and Alzheimer’s disease pathology

This TGN-retention of Sortilin is likely to have deleterious effects on many other cargoes in addition to the lysosomal enzymes we have examined here. Sortilin is a pro-neurotrophin receptor with strong links to Alzheimer’s disease (AD) (Carlo et al., 2013), having been demonstrated to bind APP (Gustafsen et al., 2013; Yang et al., 2013), BACE1 (Finan et al., 2011) as well as APOE, and also implicated in the clearance of APOE/Aβ complex in neurons (Carlo, 2013). Interestingly, Sortilin is particularly enriched in neurites and thought to promote non-amyloidogenic cleavage of APP in neurites (Gustafsen et al., 2013; Yang et al., 2013). Thus, reduced traffic of Sortilin to neurites could promote amyloidogenic APP processing in AP-4 i^3^Neurons. In addition to the possible links to AD through Sortilin, previous studies in non-neuronal cells suggest a robust and specific interaction between AP-4 and APP (Burgos et al., 2010). Depletion of AP-4 from cultured cell lines or disrupting its interaction with exogenously expressed APP has been demonstrated to increase amyloidogenic processing of APP. While this suggests a neuroprotective role for AP-4 in Alzheimer’s disease, the physiological relevance of AP-4-APP interaction in neurons has not been established. This CRISPRi based i^3^Neuron culture system could serve as an excellent model system in the future, in exploring these interactions and determining the contribution of AP-4 dependent trafficking on neuroprotection from amyloidogenic APP processing.

### AP-4 function and axonal endo-lysosome transport

In addition to changes in composition and function of lysosomes in the soma, LAMP1-positive organelles build up in axonal swellings in AP-4 i^3^Neurons with age. Since alterations to retrograde axonal transport of LAMP1-positive organelles (a mixture of late endosomes and amphisomes arising from fusion of endolysosomes with autophagosomes) have been found to cause such focal lysosomal accumulation in axonal swellings (Gowrishankar et al., 2017; Gowrishankar et al., 2021), we examined distribution of JIP3/MAPK8IP3, a potential adaptor that links axonal lysosomes to motors (Gowrishankar et al., 2017). We found that there were large accumulations of JIP3 in axonal swellings. While the build-up of axonal LAMP1-positive vesicles could trigger a compensatory increase in JIP3 levels, the accumulation of JIP3 we observed in these swellings could also result from a failure of recruitment of cytosolic JIP3 to axonal lysosomes in AP-4 i^3^Neurons (Fig. 6). *De novo* variants in *JIP3/MAPK8IP3* have been causally implicated in a neurodevelopmental disease characterized by intellectual disability, spasticity, cerebral atrophy and thin corpus callosum (Platzer et al., 2019; Iwasawa et al., 2019). Notably, these features overlap with the AP-4 deficiency syndrome, suggesting that the pathogenesis of these conditions may be linked, and the proteins may function in the same pathway.

The AP-4 complex has recently been demonstrated to interact with Hook proteins (Mattera et al., 2020) which act as dynein-dynactin activators and thus aid in retrograde transport of organelles. Indeed, through interaction with FTS (fused toes homolog) Hook and FHIP (FTS and Hook-interacting protein) proteins (FHF complex), AP-4 complex has been suggested to effect perinuclear clustering of ATG9A containing vesicles (Mattera et al., 2020). Hook proteins have been implicated in retrograde transport of BDNF-signaling endosomes in neurons (Olenick et al., 2019). The accumulation of cytosolic JIP3 in the axonal swellings in AP-4 i^3^Neurons could also be due to absence/reduced levels of interacting partners on lysosomes, whose sorting from TGN to the organelle is AP-4 dependent, as with ATG9A and potentially Sortilin. Indeed, a recent study has highlighted how TGN-derived transport carriers deliver lysosomal components to the maturing axonal organelles (Lie et al., 2021). Thus, in the absence of optimal AP-4 complex function, sorting of this putative JIP3-binding lysosomal protein into TGN-derived transport carriers would be compromised, resulting in stalled LAMP1-positive organelles in axons. Identifying AP-4 interactors in this i^3^Neuron system will shed more light on the how AP-4 regulates neuronal lysosome composition, function and transport and the mechanism underlying pathology of AP-4 deficiency syndrome.

## Material and Methods

### Generation of AP-4 CRISPRi iPSC lines

#### Constructs

Two different sgRNA sequences targeting the transcription start site of AP4E1 (G2i and G4i) were selected from the Weissman CRISPRi v2 library (Horlbeck et al., 2016). Sense and antisense sgRNA oligonucleotides were designed with 5’CACC and 3’CAAA overhangs, respectively, and cloned into pKLV-U6gRNA-EF(BbsI)-PGKpuro2ABFP for lentivirus production. Sequences of sgRNA used are detailed in Table S2. pKLV-U6gRNA-EF(BbsI)-PGKpuro2ABFP was a gift from Kosuke Yusa (Addgene plasmid # 50946; RRID: Addgene_50946).

#### Stable Cell Lines

The sgRNA lentivectors described earlier were packaged into lentivirus for transduction of iPSCs as previously described (Rodger et al., 2020). In brief, HEK293T cells were co-transfected with a lentiviral sgRNA expression plasmid and the packaging vectors pCMVΔ8.91 and pMD VSV-G (1:0.7:0.3 mix) using TransIT-293 (Mirus Bio). HEK293T media was collected 48 h post-transfection, filtered using a 0.45 μm filter, and applied to target iPSCs in the presence of 10 μg/mL polybrene (Sigma-Aldrich). Cells were transduced for 16 h and selected using 1 μg/mL puromycin 24 h later.

### iPSC culture and neuronal differentiation

iPSCs were cultured and i^3^Neurons were differentiated from them as previously described (Fernandopulle et al., 2018; Gowrishankar et al., 2021), with slight modifications. Briefly, on day 0, iPSCs were dissociated into single cells using Accutase (Thermo Fisher Scientific) and seeded at a density of 100,000 cells/well on a Matrigel-coated 6-well plate in Induction Medium composed of KO DMEM, 1× N-2 Supplement, 1× MEM Non-Essential Amino Acids Solution, 1× GlutaMAX Supplement (Thermo Fisher Scientific), 10 μM Y-27632 (Tocris Biosciences), and 2 μg/mL doxycycline hydrochloride (Sigma-Aldrich). Pre-differentiated cells were maintained in IM for 3 days with daily changes of media. After the 3-day differentiation period, cells were dissociated with Accutase and seeded at 30,000 cells per 35 mm glass bottom dish (MatTek Life Sciences) or 12mm glass coverslip (Deckglaser) coated with 0.1 mg/mL poly-L-ornithine (Sigma-Aldrich) and 10 μg/mL mouse Laminin (Gibco). Cells were maintained in Cortical Neuron Culture Medium, composed of KO DMEM (Gibco), 1× B-27 Supplement (Thermo Fisher Scientific), 10 ng/mL BDNF (PeproTech), 10 ng/mL NT-3 (PeproTech), and 1 μg/mL mouse Laminin (Thermo Fisher Scientific) with half media changes carried out every 3-4 days.

When required, as an alternative to Matrigel, vitronectin (VTN-N; Gibco Life Technologies) was used to coat the surface prior to iPSC culture as described previously (Chen et al., 2011). Briefly, VTN-N aliquots from −80°C were thawed and resuspended in Calcium- and Magnesium-free PBS to a final working concentration of 5μg/mL and used to coat the surface of 6-well plates (1 ml/well). The plates were incubated at room temperature for 1 hour and then aspirated prior to passaging the cells on to VTN-N coated plates. i^3^Neurons were plated at 35% higher density for experiments when the induction and passaging of iPSCs were on VTN-N coated plates.

### Immunofluorescence analysis of i^3^Neurons

i^3^Neurons were differentiated for one to six weeks on 35 mm glass bottom dishes (MatTek Life Sciences) and processed for immunostaining as described previously (Gowrishankar et al., 2021). See Table S1 for antibody information.

### Immunoblotting experiments

i^3^Neurons were grown on PLO/Laminin-coated 6-well plates (500,000 cells/well). After 21 days of differentiation, i^3^Neurons were washed with ice-cold PBS and lysed in Lysis buffer [Tris-buffered saline (TBS) with 1% Triton, protease inhibitor cocktail and phosphatase inhibitor] and then spun at 13,000 g for 5 minutes. The supernatant was collected and incubated at 95°C for 5 minutes in SDS sample buffer before SDS-PAGE, transfer to nitrocellulose membranes, and immunoblotting. See Table S1 for antibody information.

### Microscopy

Standard confocal images were acquired using a Zeiss 880 Airyscan confocal microscope via a 100X plan-Apochromatic objective (1.46 NA) with 2X optical zoom. Live imaging of lysosome dynamics in i^3^Neurons was carried out using Airyscan imaging mode using 100X objective with 1.5x or 2x optical zoom and scan speeds of 1-2 frames per second. Zeiss Zen software was used for processing of the Airyscan images. Further image analysis was performed using FIJI/ImageJ software (Schindelin et al., 2012).

### DQ-BSA Assay

DQ-BSA assay was carried out to examine degradative/functional lysosomes in DIV 14 i^3^Neurons grown on 35 mm glass bottom dishes (MatTek) as previously described (Marwaha and Sharma, 2017; Kulkarni et al., 2021), with minor modifications. DIV 14 i^3^Neurons were pulsed with equal concentrations of the DQ-Red BSA and Alexa 488-BSA (BSA-488) probes# (25 μg/mL, Thermo Fischer) for a period of 5 hours, followed by gentle washes [two times with warm imaging media (IM)*]. They were then imaged live in IM at 37°C using a Zeiss 880 confocal microscope in Airyscan mode with a 60x objective (NA 1.4). Healthy i^3^Neurons were identified using brightfield mode and imaged using the 561 and 488 lasers to capture fluorescence of DQ-Red BSA and BSA-488. Fluorescence intensity for each of the two channels was measured following outlining of individual cells using Image J software and the normalized ratio of DQ-Red BSA to BSA-488 intensity was computed for each cell. The mean of these per cell ratios was then computed and normalized to the value of Control i^3^Neurons to compare across multiple experiments. #DQ-Red BSA and BSA-488 probe mixture: 10% Cortical Neuron Culture Medium (Gowrishankar et al., 2021) in KO DMEM F12 was equilibrated at 37°C and 5% CO2. Probes were added to the 10% Cortical Neuronal Culture Medium and incubated at 37°C and 5% CO2 for 5 minutes for equilibration. Composition of IM: 20 mM HEPES, 5 mM KCL, 1 mM CaCl2, 150 mM NaCl, 1 mM MgCl2, 1.9 mg/mL glucose and BSA, pH 7.4. The probe mixture pulses were staggered by 45 minutes between dishes to enable live imaging.

### Analysis of vesicle count in i^3^Neurons

Confocal (LAMP1, Cathepsin B and L, PPT-1) and Airy scan images (DQ-BSA) were used for analysis of vesicle size and number in i^3^Neurons. Vesicles were identified using “threshold” function and then their number and size determined using “Analyze particles” function in Fiji/ImageJ software. Detailed statistics are included in the figure legend.

### Analysis of axonal accumulation of LAMP1 and JIP3

Maximum intensity projections of confocal images obtained of LAMP1 or JIP3 staining in i^3^Neurons were used for analyzing the extent of axonal accumulation of these proteins. Images from Control and AP-4 i^3^Neurons were thresholded to the same intensity values and accumulations in the axons identified as those larger than 2 μm and above the threshold. All such “accumulations” were counted irrespective of relative size and intensity. The axonal accumulations were reported as the number per 18,211 μm^2^ (area of each confocal image).

### Colocalization analysis of lysosomal enzymes with LAMP1

Maximum intensity projections of confocal images obtained of LAMP1 co-stained with Cathepsin L or PPT-1 in i^3^Neurons were used for analyzing the fraction of each of those enzymes that overlaps with LAMP1. The images are thresholded and the extent/fraction of PPT-1/Cathepsin L that overlapped with LAMP1 (i.e., overlap of PPT-1 or Cathepsin L channel with the LAMP1 channel) was computed for each soma, using JACOP plugin of Image J.

### Statistical analysis

Data are represented as mean ± SEM unless otherwise specified. Statistical analysis was performed using Prism 8 software. Groups were compared using one-way ANOVA with Dunnett’s multiple comparison test. Detailed statistical information (number of independent experiments, and p-values) is described in the respective Figure legends.

## Acknowledgement

We thank Daniel McBride an Eduardo Pallares for technical assistance. We thank Dr. Hofmann (UT Southwestern) for generous gift of the PPT-1 antibody. We thank Michael Ward (NIH) for the generous gift of CRISPRi – i^3^ neurons. SG is funded by the Wolverine foundation and the Dr. Ralph and Marian Falk Medical Research Trust. ER was funded by the NIHR Cambridge Biomedical Research Centre, the UK Medical Research Council (Project grant MR/R026440/1) and a generous donation from Hazel and Keith Satchell. The views expressed are those of the authors and not necessarily those of the NIHR or the UK Department of Health and Social Care. The authors declare no competing financial interests.

## Author Contributions

PM helped write the manuscript and led efforts related to experiment design, performing experiments, data analysis, and figure preparation. DE performed experiments, analyzed data and helped with manuscript editing. CR generated CRISPRi iPSC lines and helped with manuscript editing. SP performed experiments and analyzed data. ER supervised generation of iPSC lines, contributed to experimental design and manuscript writing. SG supervised the project, designed, and performed experiments, analyzed the data, and led the writing of the manuscript. Through ongoing discussion of results, all authors contributed to the overall direction of the project.

