## Supplemental Figures and Legends and Supplemental Tables for "AP-4 regulates neuronal lysosome composition, function, and transport via regulating export of critical lysosome receptor proteins at the trans-Golgi network"

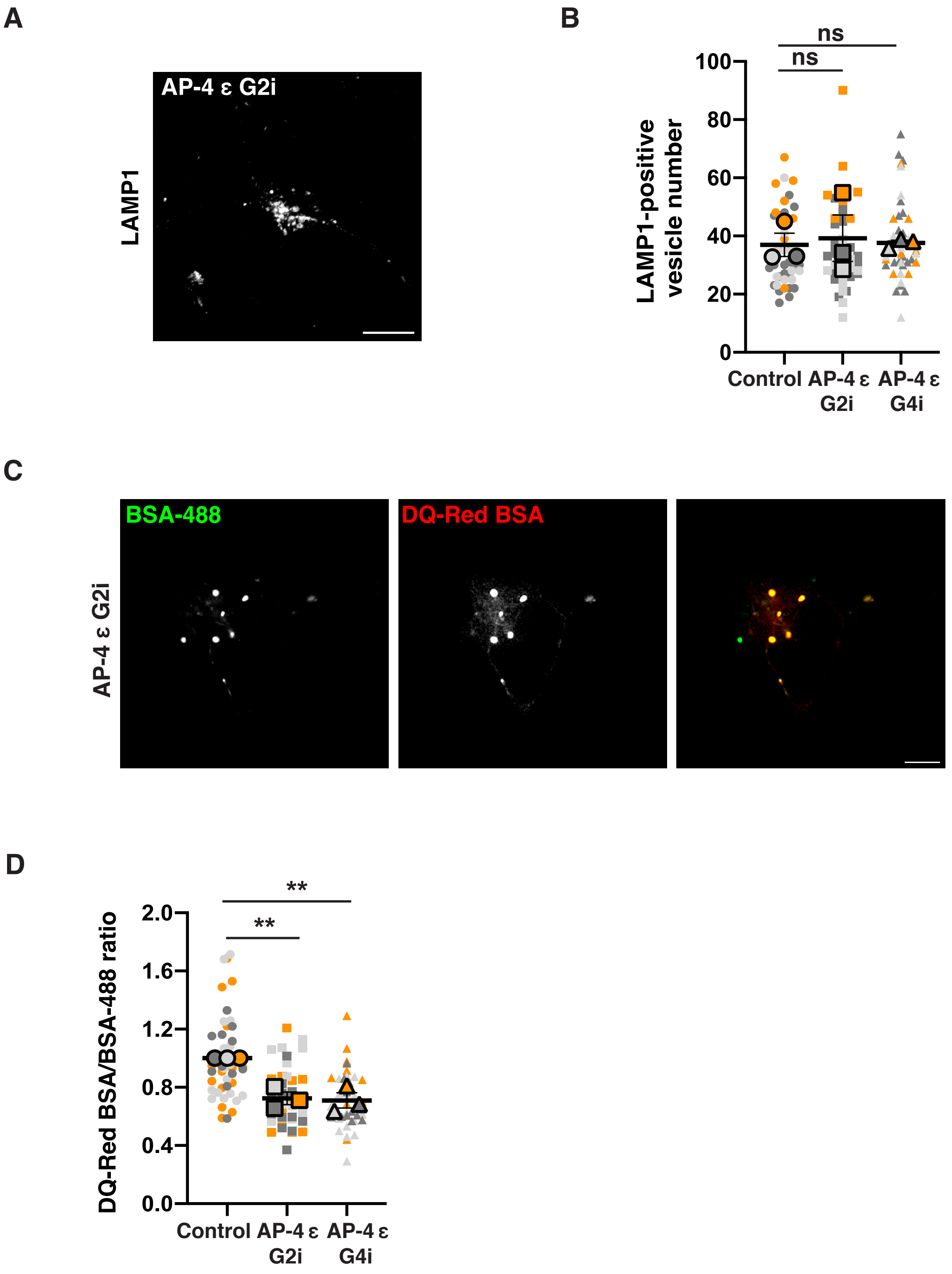
A
AP-4 ε G2i
LAMP1
B
ns
ns
LAMP1-positive vesicle number
100
80
60
40
20
0
Control
AP-4 ε G2i
AP-4 ε G4i
C
BSA-488
DQ-Red BSA
AP-4 ε G2i
D
**
**
DQ-Red BSA/BSA-488 ratio
2.0
1.6
1.2
0.8
0.4
0.0
Control
AP-4 ε G2i
AP-4 ε G4i

Figure S1

**Figure S1. Loss of AP-4 affects neuronal lysosome function.**

(A) AP-4 G2i $i^3$Neurons (14 DIV) stained for LAMP1 compared to its corresponding Control in Fig 1C. Bar, 10 µm. (B) Superplot depicting the number of LAMP1-positive vesicles in soma of each $i^3$Neuron in all the genotypes (SC-circle, AP-4 G2i-square, and AP-4 G4i-triangle) from three independent experiments (experiments depicted in steel, silver and tangerine colors), as well as the population mean of each experiment (larger symbols with bold outline) and SEM. Superplot thus depicts cell to cell and experimental variability. (C) Images of DQ-Red BSA and A488 BSA fluorescence in AP-4 G2i $i^3$Neuron compared to its corresponding Control in Fig 1E (14 DIV). Bar, 10 µm. (D) Superplot depicting the lysosomal degradation (DQ-Red BSA/BSA-488 ratio) in each $i^3$Neuron in all the genotypes (SC-circle, AP-4 G2i-square, and AP-4 G4i-triangle) across three independent experiments, where ratio from each cell was in turn normalized to population mean of SC $i^3$Neurons from same experiment. Superplot also depicts population mean of each experiment (larger symbols with bold outline) and SEM.

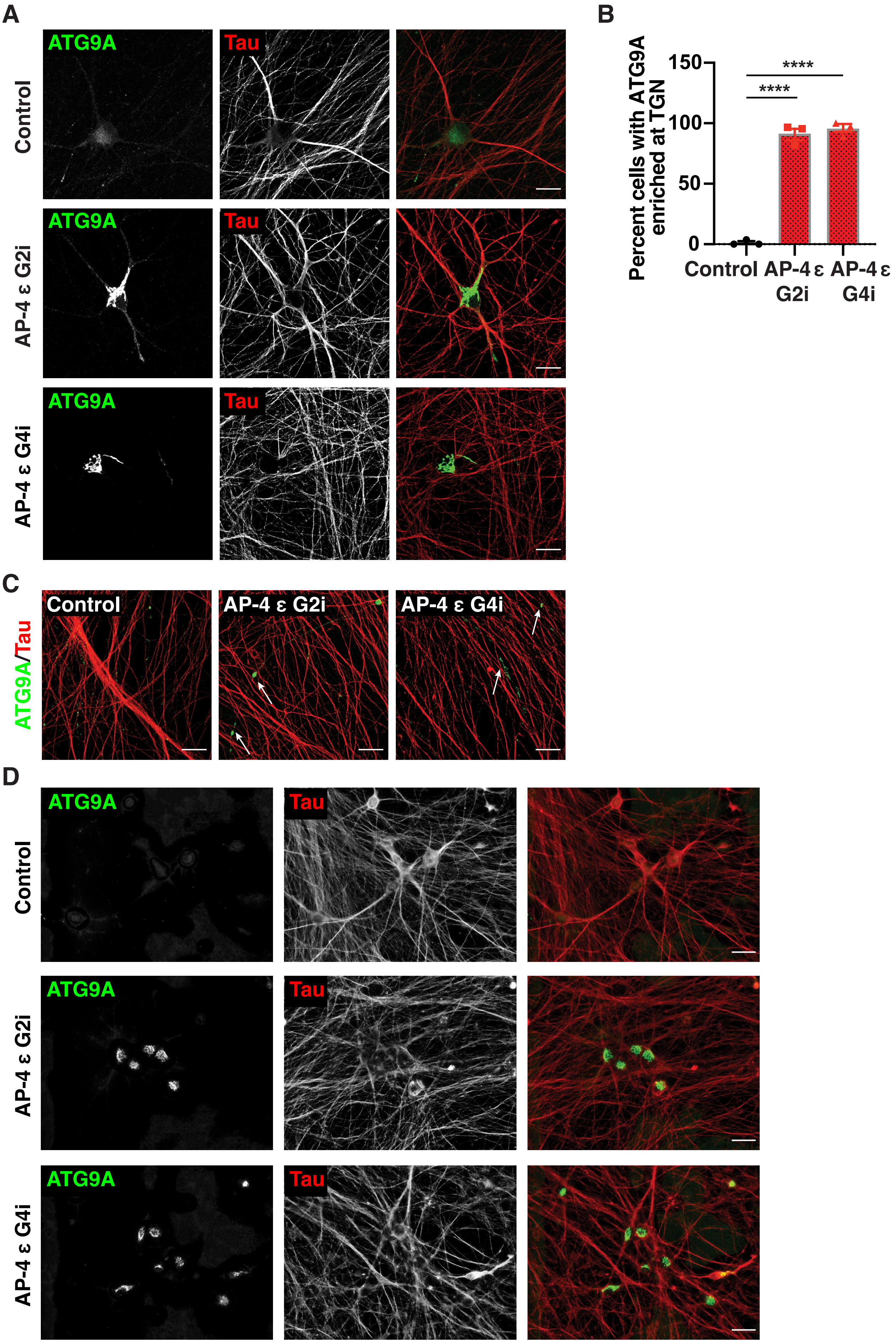
A
ATG9A
Tau
Control
AP-4 ε G2i
AP-4 ε G4i
B
Percent cells with ATG9A enriched at TGN
150
100
50
0
****
****
Control
AP-4 ε G2i
AP-4 ε G4i
C
ATG9A/Tau
D

Figure S2

**Figure S2. ATG9A is strongly retained in TGN area in AP-4 i$^3$Neurons.**

(A) High-resolution confocal images of Control and AP-4 i$^3$Neurons (42 DIV) stained for ATG9A (green) and Tau (red) showing strong accumulation of ATG9A in the TGN area of the soma in AP-4 i$^3$Neurons. Bar, 20 µm. (B) Quantification of the percentage of neurons with ATG9A accumulating in the soma in AP-4 i$^3$Neurons compared to Control i$^3$Neurons (42 DIV; mean ± SEM from three independent experiments; >75 neurons per genotype; ****, $P < 0.0001$). (C) Representative images of Control and AP-4 i$^3$Neurons (42 DIV) stained for ATG9A (green) and Tau (red). Arrows highlight ATG9A accumulation in the neurites of AP-4 i$^3$Neurons. Bar, 20 µm. (D) Representative epifluorescence images showing immunofluorescence of ATG9A (green) and Tau (red) in multiple Control and AP-4 i$^3$Neurons (42 DIV) within a single frame. Bar, 50 µm.

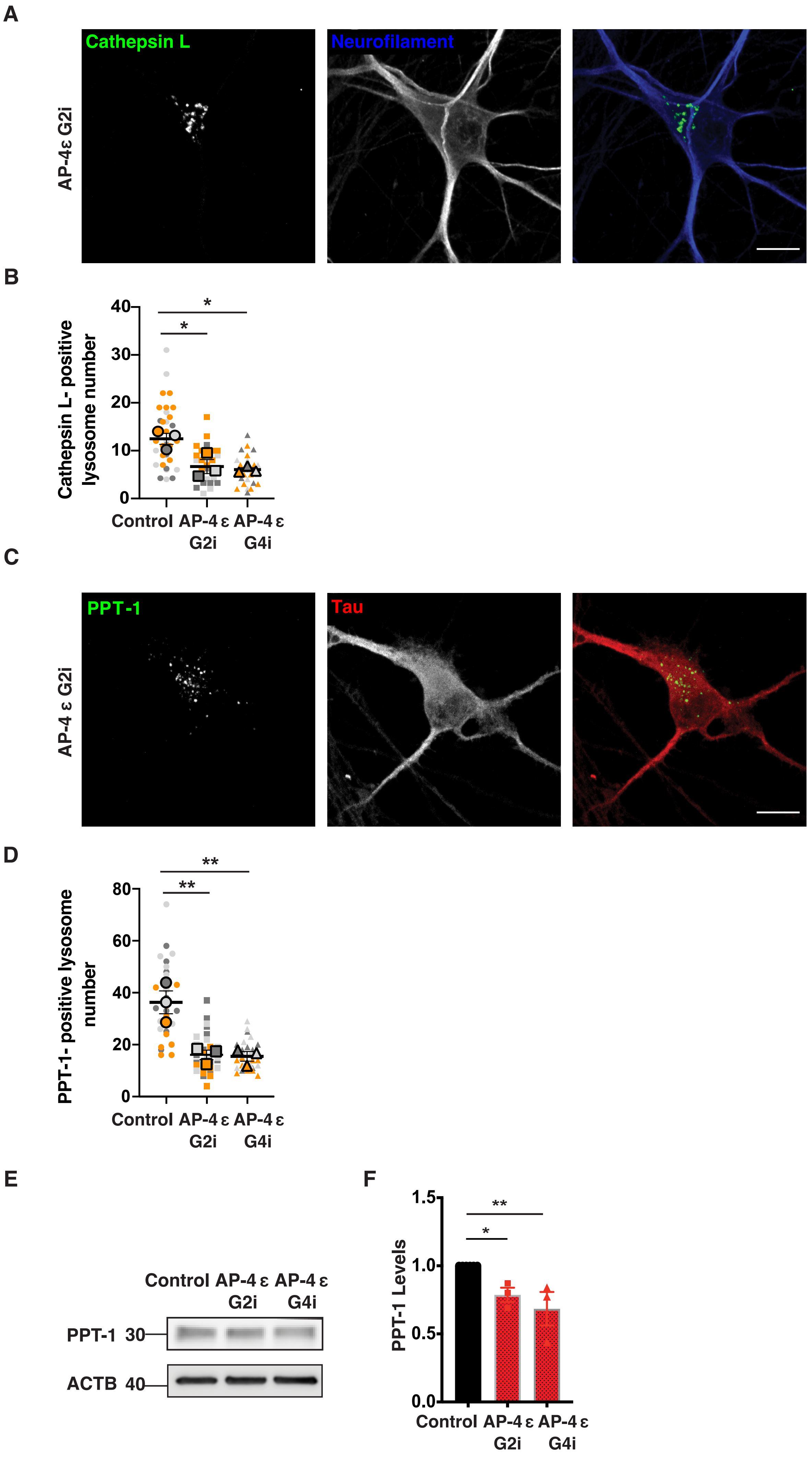
A
Cathepsin L
Neurofilament
AP-4ε G2i
B
Cathepsin L- positive lysosome number
Control
AP-4 ε G2i
AP-4 ε G4i
C
PPT-1
Tau
AP-4 ε G2i
D
PPT-1- positive lysosome number
E
Control AP-4 ε G2i AP-4 ε G4i
PPT-1 30
ACTB 40
F
PPT-1 Levels

Figure S3

**Figure S3. Loss of AP-4 affects neuronal lysosomal composition.**

(A) AP-4 G2i i$^3$Neuron (14 DIV) stained for Cathepsin L (green) and neurofilament (blue) showing relatively reduced number of vesicles compared to its corresponding Control in Fig 2C. Bar, 10 µm. (B) Superplot depicting the number of Cathepsin L-positive lysosomes in soma of each i$^3$Neuron in all the genotypes (SC-circle, AP-4 G2i-square, and AP-4 G4i-triangle) from three independent experiments (experiments depicted in steel, silver and tangerine colors), as well as the population mean of each experiment (larger symbols with bold outline) and SEM. (C) AP-4 G2i i$^3$Neurons (14 DIV) stained for PPT-1 (green) and Tau (red) showing relatively reduced number of PPT-1 vesicles compared to its corresponding Control in Fig 2E. Bar, 10 µm. (D) Superplot depicting the number of PPT-1-positive lysosomes in soma of each i$^3$Neuron in all the genotypes (SC-circle, AP-4 G2i-square, and AP-4 G4i-triangle) from three independent experiments (experiments depicted in steel, silver and tangerine colors), as well as the population mean of each experiment (larger symbols with bold outline) and SEM. (E and F) Immunoblotting reveals decreased levels of PPT-1 protein in AP-4 i$^3$Neurons compared to Control i$^3$Neurons (21 DIV; ACTB used as loading Control; mean ± SEM from three independent experiments; *, $P < 0.05$; **, $P < 0.01$).

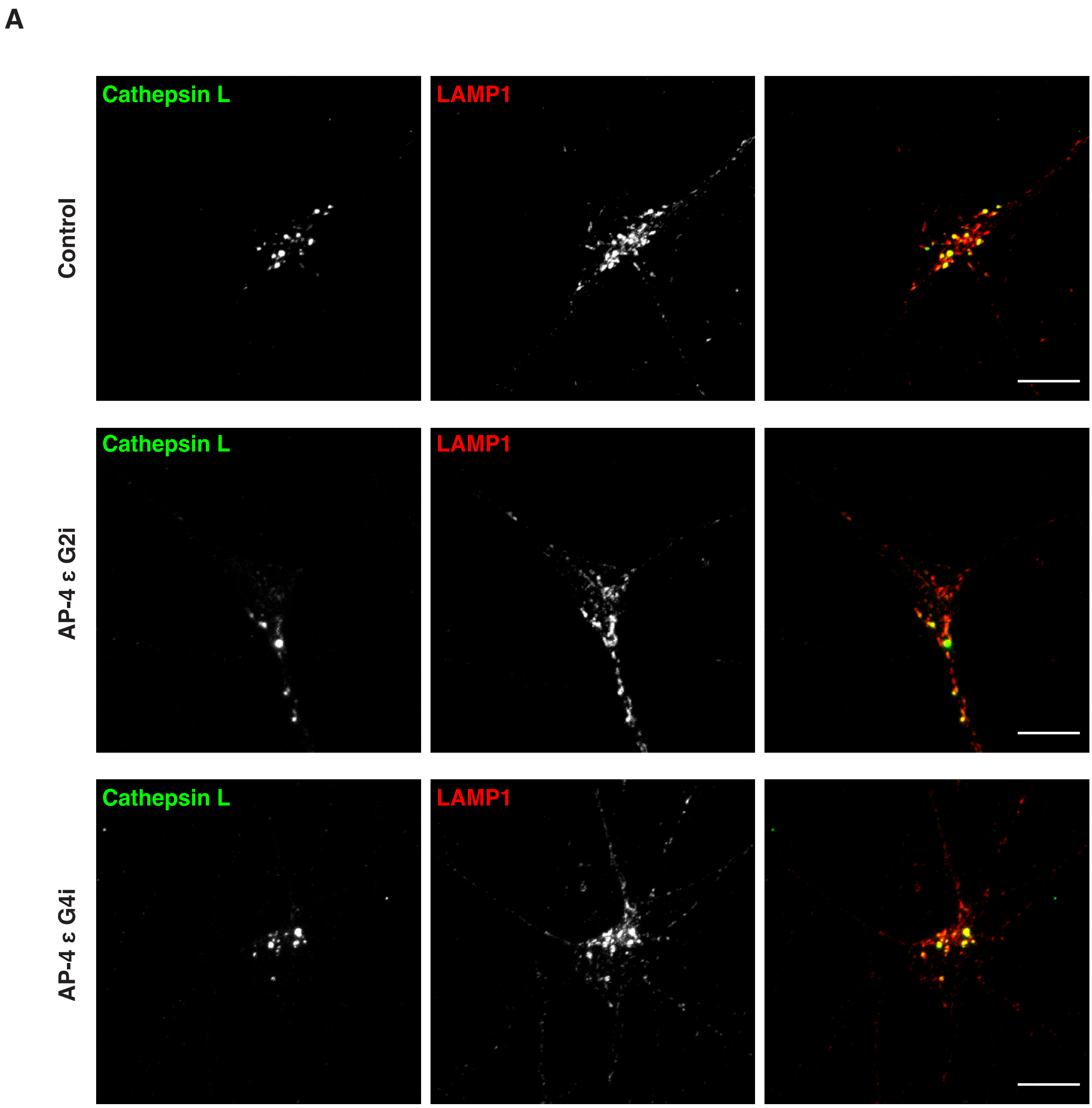
A
Cathepsin L
LAMP1
Control
Cathepsin L
LAMP1
AP-4 ε G2i
Cathepsin L
LAMP1
AP-4 ε G4i

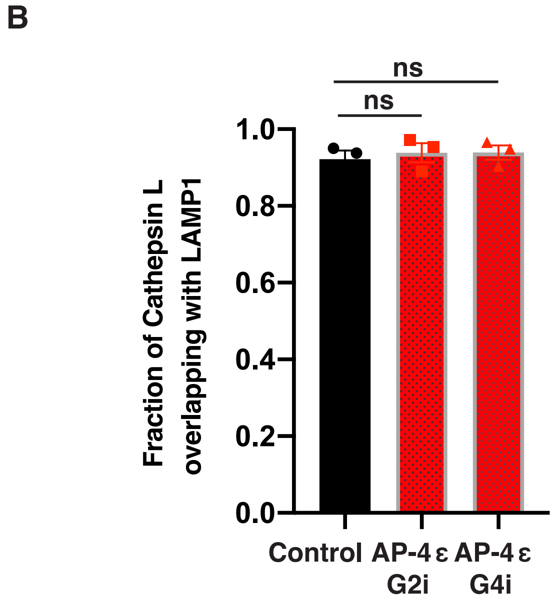
B
ns
ns
1.0
0.8
0.6
0.4
0.2
0.0
Fraction of Cathepsin L overlapping with LAMP1
Control
AP-4 ε G2i
AP-4 ε G4i

Figure S4

**Figure S4. Cathepsin L-enriched vesicles are LAMP1 positive in i$^3$Neurons of all genotypes**

(A) Control and AP-4 i$^3$Neurons (14 DIV) co-stained for Cathepsin L and LAMP1 reveals that the Cathepsin L-positive vesicles are enriched for LAMP1 in all genotypes, even though AP-4 i$^3$Neurons have fewer Cathepsin L-positive vesicles. Bar, 10 µm. (B) Quantification of the colocalization of Cathepsin L with LAMP1 (examining fraction of Cathepsin L channel that overlaps with LAMP1), reveals that the fraction of Cathepsin L that overlaps with LAMP1 is not altered in AP-4 i$^3$Neurons compared to Control i$^3$Neurons, suggesting that all the detected Cathepsin L vesicles represent lysosomal signal (14 DIV; mean ± SEM from three independent experiments; 30-35 per genotype; ns, not significant).

A

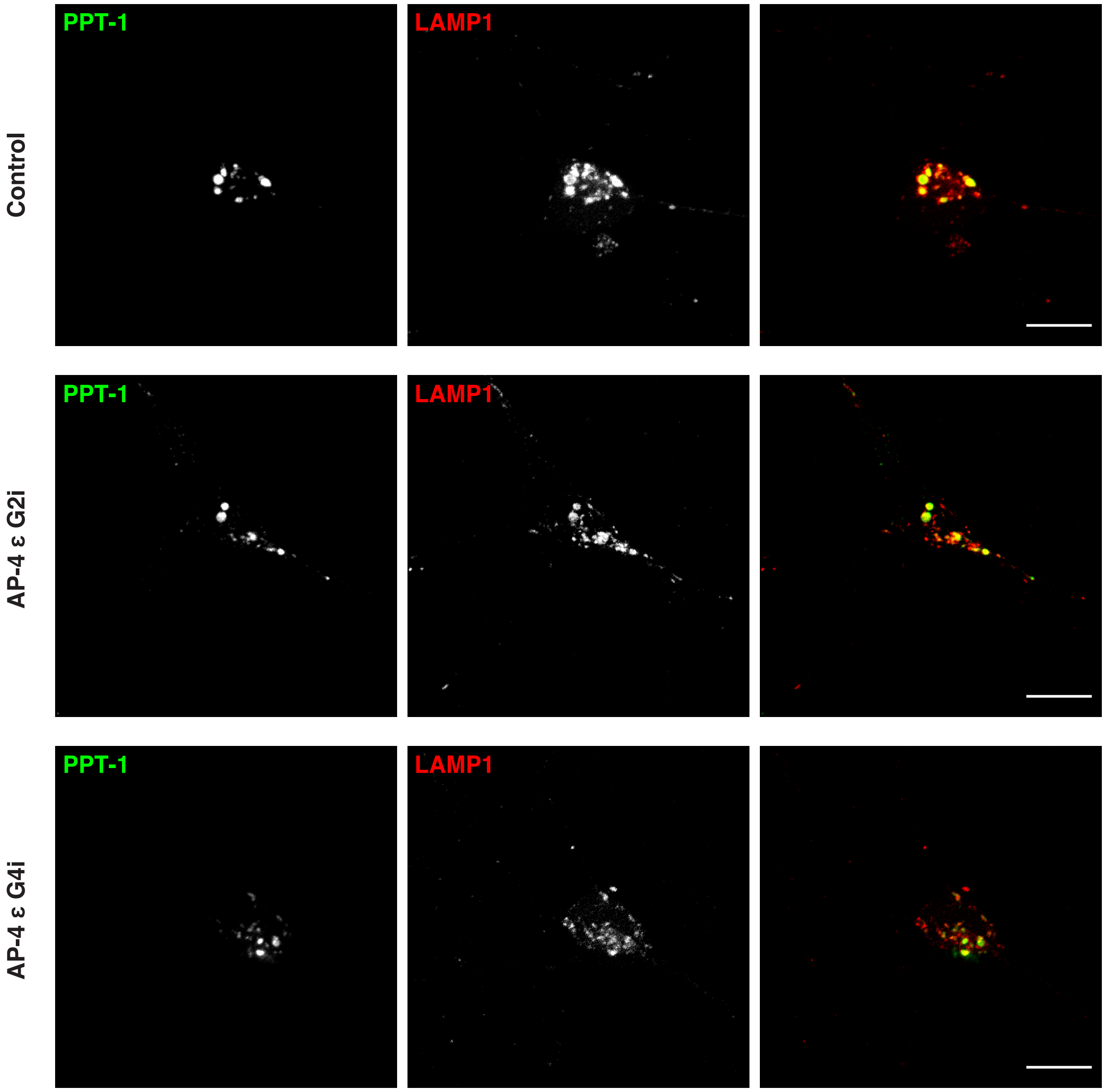

PPT-1
LAMP1
Control
PPT-1
LAMP1
AP-4 ε G2i
PPT-1
LAMP1
AP-4 ε G4i


B

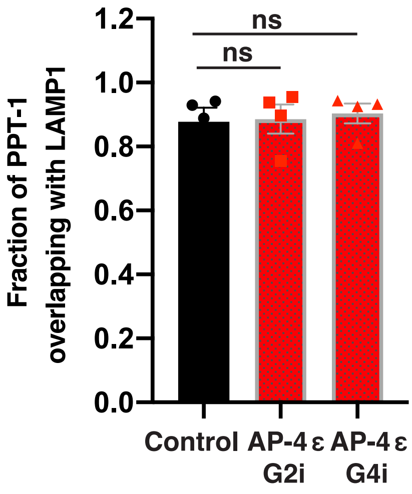

Fraction of PPT-1 overlapping with LAMP1
1.2
1.0
0.8
0.6
0.4
0.2
0.0
ns
ns
Control
AP-4 ε G2i
AP-4 ε G4i


Figure S5

**Figure S5. PPT-1-enriched vesicles are LAMP1 positive in i$^3$Neurons of all genotypes**

(A) Control and AP-4 i$^3$Neurons (14 DIV) co-stained for PPT-1 and LAMP1 reveals that the PPT-1-positive vesicles are enriched for LAMP1 in all genotypes, even though AP-4 i$^3$Neurons have fewer PPT-1-positive vesicles. Bar, 10 µm. (B) Quantification of the colocalization of PPT-1 with LAMP1 (examining fraction of PPT-1 channel that overlaps with LAMP1), reveals that the fraction of PPT-1 that overlaps with LAMP1 is not altered in AP-4 i$^3$Neurons compared to Control i$^3$Neurons, suggesting that all the detected PPT-1 vesicles represent lysosomal signal (14 DIV; mean ± SEM from four independent experiments; 30-35 per genotype; ns, not significant).

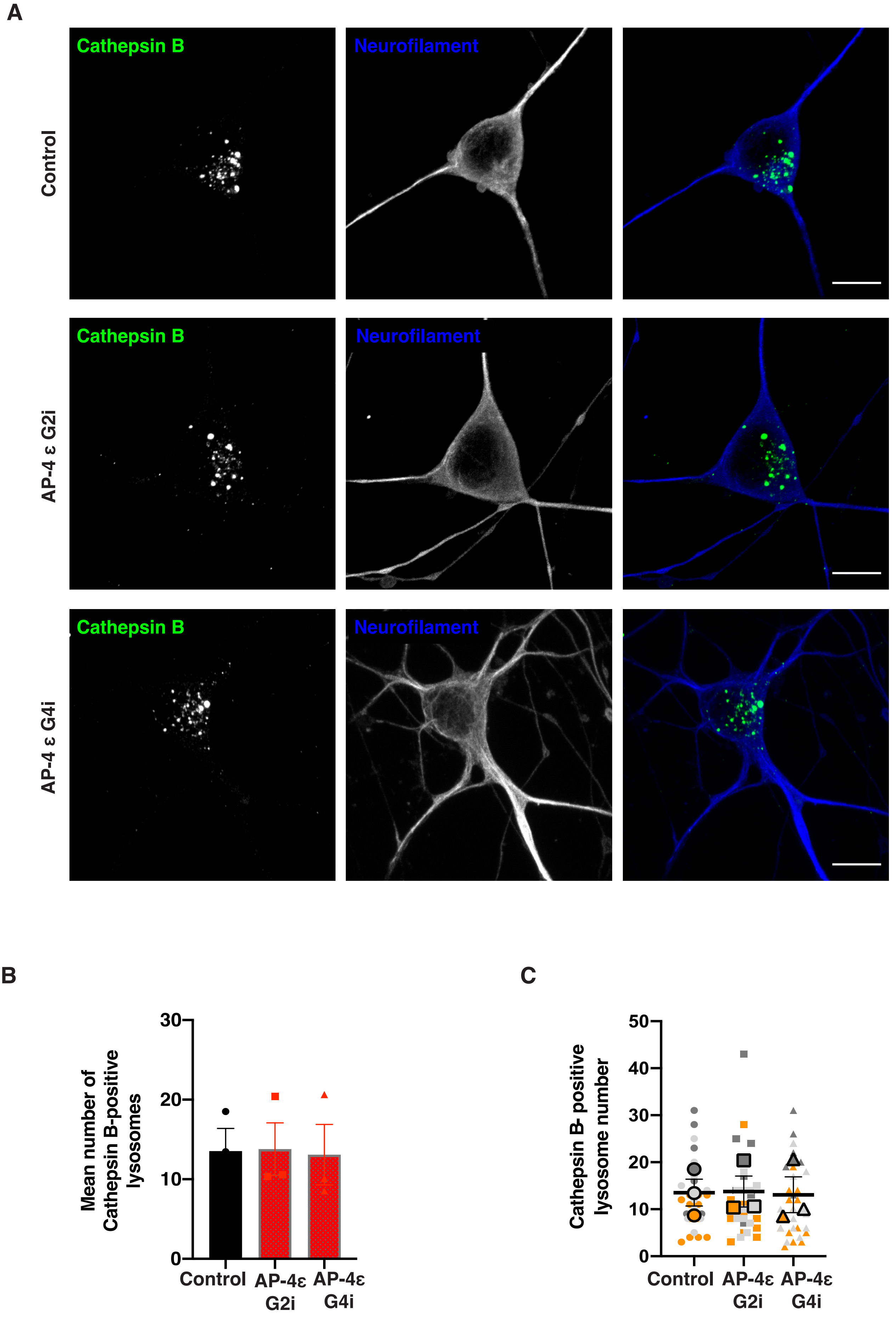

A
Cathepsin B
Neurofilament
Control
Cathepsin B
Neurofilament
AP-4 ε G2i
Cathepsin B
Neurofilament
AP-4 ε G4i
B
Mean number of Cathepsin B-positive lysosomes
30
20
10
0
Control
AP-4ε G2i
AP-4ε G4i
C
Cathepsin B- positive lysosome number
50
40
30
20
10
0
Control
AP-4ε G2i
AP-4ε G4i


Figure S6

**Figure S6. Loss of AP-4 does not affect Cathepsin B traffic to lysosomes.**

(A) Control and AP-4 $i^3$Neurons (14 DIV) stained for Cathepsin B (green) and neurofilament (blue). Bar, 10 µm. (B) Quantification of mean number of Cathepsin B-positive lysosomes in AP-4 $i^3$Neurons compared to Control $i^3$Neurons (14 DIV; mean ± SEM from three independent experiments; >30 neurons per genotype; ns, not significant). (C) Superplot depicting the number of Cathepsin B-positive lysosomes in soma of each $i^3$Neuron in all the genotypes (SC-circle, AP-4 G2i-square, and AP-4 G4i-triangle) from three independent experiments (experiments depicted in steel, silver and tangerine colors), as well as the population mean of each experiment (larger symbols with bold outline) and SEM.

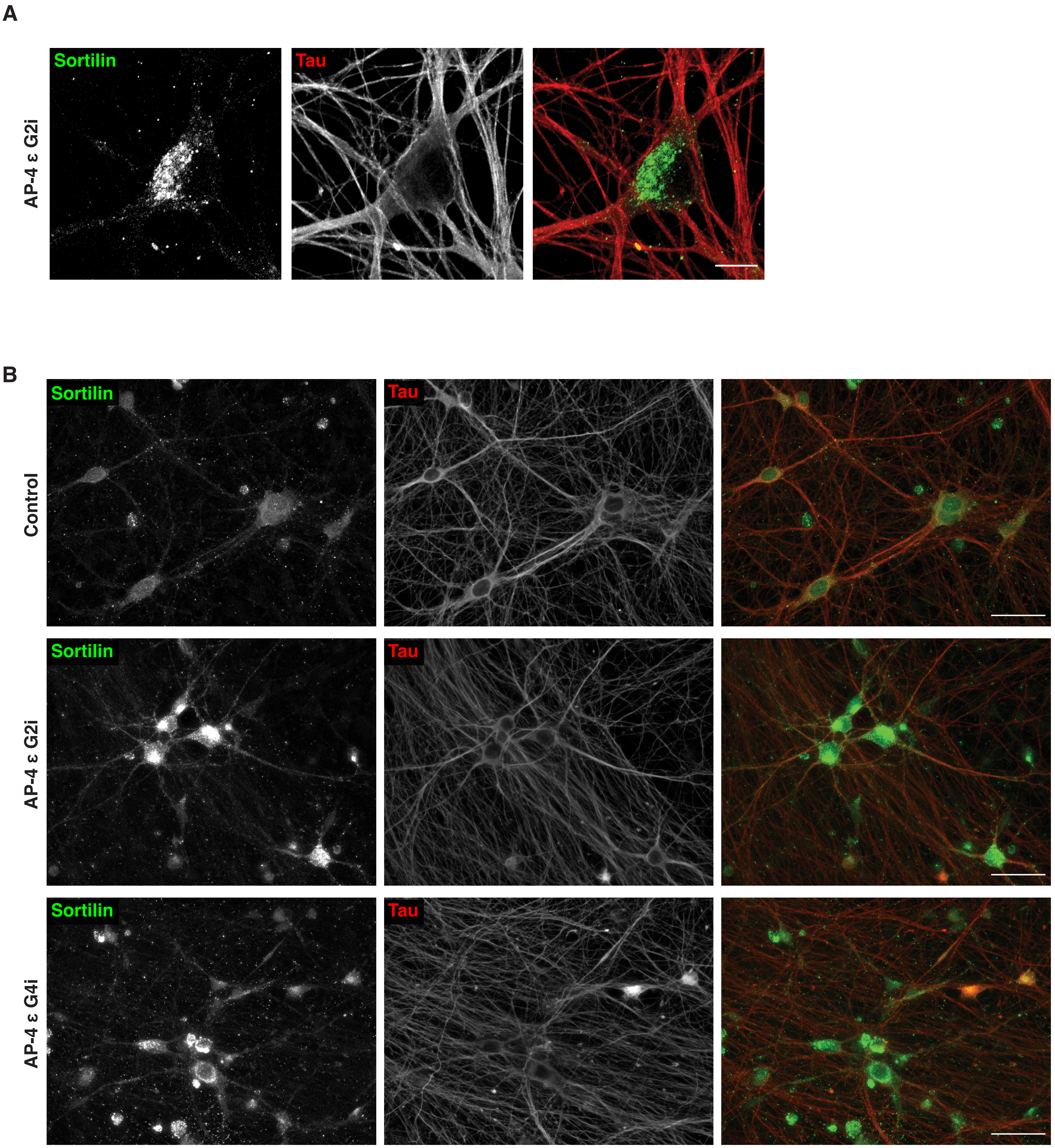
A
Sortilin
Tau
AP-4 ε G2i
B
Sortilin
Tau
Control
Sortilin
Tau
AP-4 ε G2i
Sortilin
Tau
AP-4 ε G4i

**Figure S7**

**Figure S7. Sortilin distribution is dramatically altered in AP-4 i$^3$Neurons.**

(A) High resolution images for AP-4 G2i i$^3$Neurons (42 DIV) stained for Sortilin (green) and Tau (red), showing strong enrichment in the TGN area compared to its corresponding Control in Fig 3C. Bar, 10 µm. (B) Representative epifluorescence images showing immunofluorescence of Sortilin (green) and Tau (red) in multiple Control and AP-4 i$^3$Neurons (42 DIV) within a single frame. Bar, 50 µm.

A

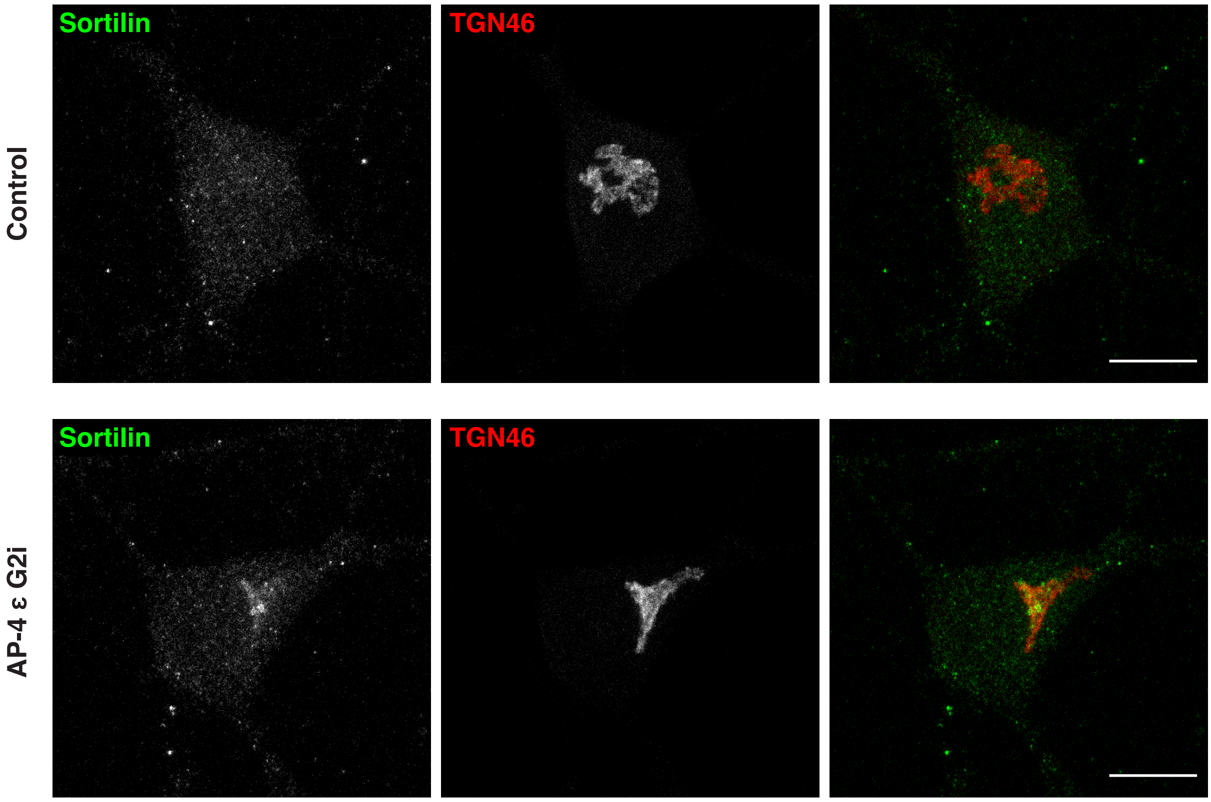


Figure S8

**Figure S8. Sortilin accumulating in soma of AP-4 i$^3$Neurons colocalizes with the Trans Golgi**

(A) High resolution images of Control and AP-4 G2i i$^3$Neuron (28 DIV) stained for Sortilin (green) and TGN46 (red) reveals that Sortilin accumulation colocalizes with TGN46 and is thus in the TGN of AP-4 G2i i$^3$Neurons, unlike in Control i$^3$Neurons where there is no colocalization/enrichment with TGN46. Bar, 10 μm.

A

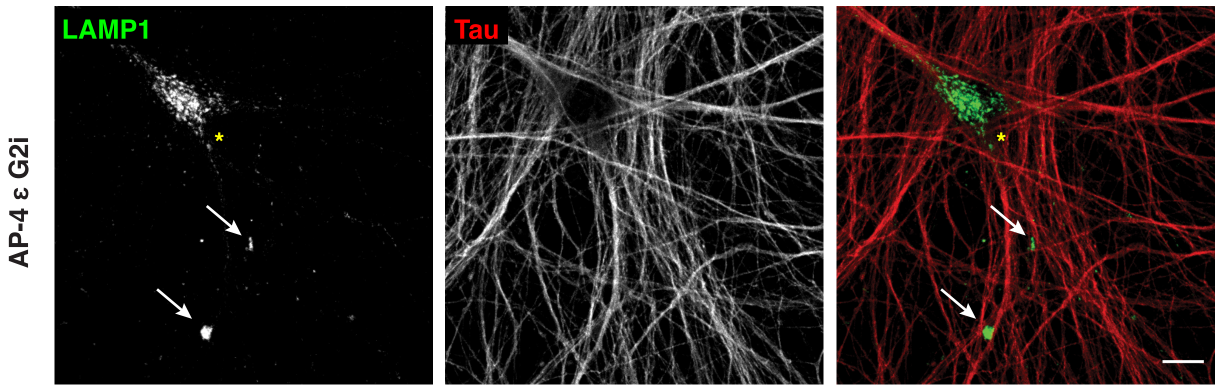


B

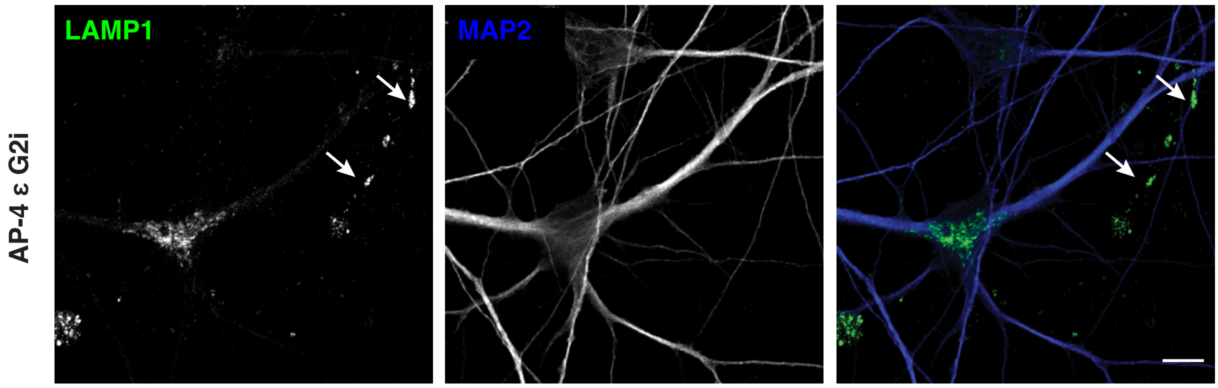


C

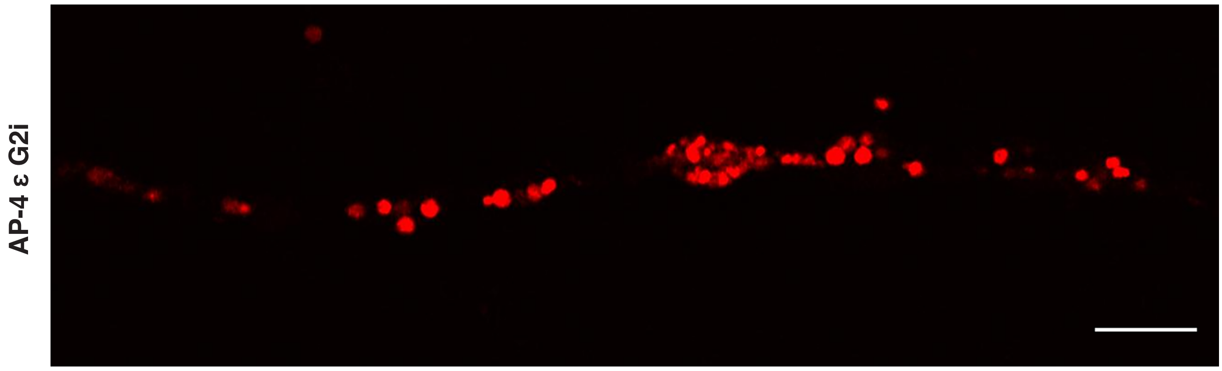


D

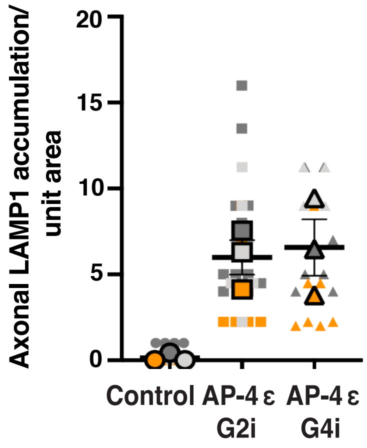


Figure S9

**Figure S9. Axonal phenotypes in AP-4 i$^3$Neurons.**

(A) AP-4 G2i i$^3$Neurons (42 DIV) stained for LAMP1 (green) and Tau (red) compared to its corresponding Control in Fig 4A. Arrows highlight axonal lysosome accumulations in AP-4 i$^3$Neurons. Yellow asterisk indicates the lysosomes in the cell body. Bar, 20 µm. (B) AP-4 G2i i$^3$Neurons (42 DIV) stained for LAMP1 (green) and MAP2 (blue) compared to its corresponding Control in 4C. Arrows highlight LAMP1 accumulation in the MAP2-negative neurites (axons) of the AP-4 i$^3$Neurons. Bar, 20 µm. (C) LysoTracker staining in "straightened" axon of AP-4 i$^3$Neurons showing accumulation of large, acidic vesicles in axonal swelling and along the axon. Bar, 5 µm. (D) Superplot depicting the number of LAMP1-positive axonal accumulations per unit area in all the genotypes (SC-circle, AP-4 G2i-square, and AP-4 G4i-triangle) from three independent experiments (experiments depicted in steel, silver and tangerine colors), as well as the population mean of each experiment (larger symbols with bold outline) and SEM.

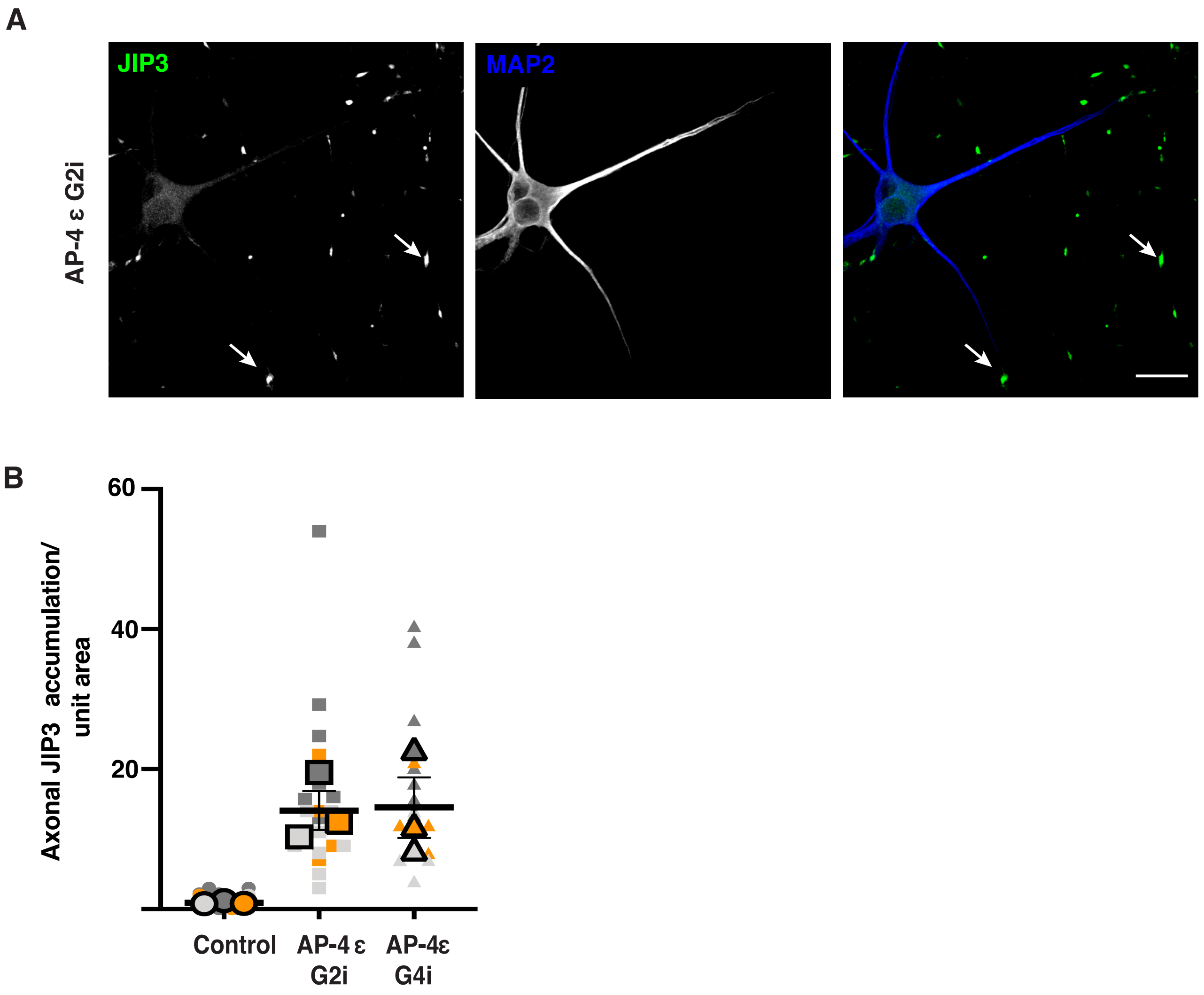
A
AP-4 ε G2i
JIP3
MAP2
B
Axonal JIP3 accumulation/ unit area
60
40
20
Control
AP-4 ε G2i
AP-4ε G4i

**Figure S10. Axonal phenotypes in AP-4 i³Neurons.**

(A) AP-4 G2i i3Neurons (28 DIV) stained for JIP3 (green) and MAP2 (blue) compared to its corresponding Control in Fig 5C. Arrows highlight JIP3 accumulation in the neurites of the AP-4 i3Neurons. Bar, 20 µm. (B) Superplot depicting the number of JIP3-positive axonal accumulations per unit area in all the genotypes (SC-circle, AP-4 G2i-square, and AP-4 G4i-triangle) from three independent experiments (experiments depicted in steel, silver and tangerine colors), as well as the population mean of each experiment (larger symbols with bold outline) and SEM.

**Table S1: Antibody list**

| Antibody | Dilution | Experiment | Company (Catalog No.) |
|---|---|---|---|
| AP-4 ε | 1:1000 | Western Blot | BD Transduction Laboratories (612018) |
| ATG9A | 1:200 | IF | Abcam (ab108338) |
| LAMP1 | 1:400 | IF | CST (9091) |
| Neurofilament | 1:400 | IF | BioLegend (837801 (SMI 311-R) |
| Cathepsin B | 1:200 | IF | R&D systems (AF953) |
| Cathepsin L | 1:1000 | Western Blot | Sigma Aldrich (C4618) |
| | 1:200 | IF | R&D systems (AF952) |
| | 1:200 | IF | |
| Tau | 1:400 | IF | CST (4019S) |
| JIP3 | 1:1000 | Western Blot | Sigma Aldrich (HPA069311) |
| | 1:200 | IF | |
| MAP2 | 1:500 | IF | Millipore (AB5622) |
| | 1:200 | | Invitrogen (PA1-10005) |
| PPT-1 | 1:1000 | Western Blot | Dr. Hofmann |
| | 1:100 | IF | |
| Sortilin | 1:100 | IF | Abcam (ab188586) |

**Table S2: Sequence of sgRNA**

| sgRNA | Sequence |
|---|---|
| Scrambled (Control) | GGGACGCGAAAGAAACCAGT |
| AP4ε1 G2 (G2i) | AAACAGGAAGTGCCTACGG |
| AP4ε1 G4 (G4i) | GGCATGAAGCCGGGCGGCTA |

**Table S3: Abbreviations**

| Abbreviation | Denotation |
|---|---|
| AD | Alzheimer’s disease |
| AP-4 | Adaptor protein complex-4 |
| APOE | Apolipoprotein E |
| APP | Amyloid precursor protein |
| ATG9A | Autophagy-related protein 9 |
| BACE1 | Beta-site APP cleaving enzyme 1 |
| BDNF | Brain-derived neurotrophic factor |
| BSA | Bovine serum albumin |
| CD63 | Cluster of differentiation 63 (Tetraspanin/LAMP3) |
| CRISPRi | CRISPR-inhibition |
| DIV | Days *in vitro* |
| FHIP Protein | FTS and Hook-interacting protein |
| FTS Hook | Fused toes homolog |
| HSP | Hereditary spastic paraplegia |
| $i^3$Neurons | iPSC-derived Glutamatergic cortical neurons |
| iPSC | Induced pluripotent stem cells |
| KRAB | Krüppel-associated box |
| LAMP1 | Lysosome associated Membrane Protein1 |
| LAMP2 | Lysosome associated Membrane Protein2 |
| M6PR | Mannose-6-phosphate receptor |
| MAP2B | Microtubule-associated protein 2 |
| MAPK8IP3/JIP3 | Mitogen-Activated Protein Kinase 8 Interacting Protein 3/ JNK-interacting protein 3 |
| MEF | mouse embryonic fibroblast |
| NGN2 | Neurogenin 2 |
| PPT-1 | palmitoyl-protein thioesterase 1 |
| sgRNA | single guide RNA |
| TGN | trans-Golgi network |

**Supplemental Video Legends**

**Endo-lysosomes accumulate in axonal swellings in AP-4 i$^3$Neurons**

Video 1: 3D reconstruction of Airyscan image of an axonal swelling filled with LAMP1-positive vesicles in AP-4 i$^3$Neurons. Video 2: Large number of LysoTracker-positive vesicles (acidic) in both the swelling and rest of the axon in an AP-4 i$^3$Neuron. Images were acquired at 1 frame/s, playback rate 7 frames/s. Scale bar, 5 µm.
